# Proximity labeling at non-centrosomal microtubule-organizing centers reveals VAB-10B and WDR-62 as distinct microtubule regulators

**DOI:** 10.1101/2020.08.29.272369

**Authors:** Ariana D. Sanchez, Tess C. Branon, Lauren E. Cote, Alexandros Papagiannakis, Xing Liang, Melissa A. Pickett, Kang Shen, Christine Jacobs-Wagner, Alice Y. Ting, Jessica L. Feldman

**Author notes:** Corresponding author: Jessica L. Feldman, Department of Biology, Stanford University, 371 Serra Mall, 94305 Stanford, USA.

## Abstract

Reorganization of microtubules from the centrosome to non-centrosomal subcellular sites is central to cell differentiation. To identify components of non-centrosomal microtubule organizing centers in differentiated cells of a living organism, we developed the biotin ligase-based proximity labeling approach TurboID for use in *C. elegans*. We identified proteins proximal to the non-centrosomal microtubule minus end protein PTRN-1/Patronin at the apical membrane of epithelial cells, focusing on two conserved proteins: spectraplakin protein VAB-10B and WDR-62, a protein we identify as homologous to vertebrate primary microcephaly disease gene WDR62. We found that WDR-62 and VAB-10B independently regulate the growth and localization of non-centrosomal microtubules and the apical targeting of microtubule minus end proteins. This regulation occurs downstream of cell polarity and in conjunction with actin. Our data suggest a division of labor where microtubule growth and anchoring are regulated by distinct complexes and uncover novel functions of spectraplakins and WDR62 family proteins.

## Introduction

Microtubules are polarized polymers critical for eukaryotic cell functions including intracellular transport, organelle positioning, cell polarity, and cell shape. Early electron microscopy studies identified subcellular sites to which microtubules localize, termed microtubule-organizing centers (MTOCs) (Porter 1966; Pickett-Heaps 1969). MTOCs are broadly defined as sites that grow and organize microtubules from their minus ends, greatly contributing to minus end stability and to the ability of plus ends to dynamically probe their environment and build specific spatial patterns. The best-studied MTOC is the centrosome, a non-membrane bound organelle that patterns microtubules into a radial array in animal mitosis and in some specialized cell types. However, in many types of differentiated cells, microtubules no longer associate with the centrosome but rather a non-centrosomal MTOC (ncMTOC) to achieve cell-type specific organization. For example, intestinal and tracheal epithelial cells organize microtubules at the apical membrane into parallel arrays along the apicobasal axis, germ cells organize microtubules at their plasma membrane, neurons organize longitudinal microtubule arrays down axons and dendrites in some cases around endosomes, and muscle cells can organize microtubules from Golgi outposts or the nuclear envelope (Brodu et al., 2010; Yang & Feldman 2015; Zhou et al., 2009; Oddoux et al., 2013; Ori-McKenney et al., 2012; Liang et al. 2020). Despite the ubiquity of ncMTOCs across cell types and organisms, their molecular components and their mechanisms of formation remain largely unknown.

The ability of MTOCs to grow, stabilize, and anchor microtubules is imparted by molecules acting alone or in functional complexes at microtubule minus ends, but our knowledge of minus end proteins or how they are targeted to specific cellular sites is scarce. The first identified microtubule minus end complex, the γ-tubulin ring complex (γ-TuRC), functions as a microtubule nucleator and minus end stabilizer (C. E. Oakley & B. R. Oakley 1989; Wiese & Zheng 2000). γ-TuRC localizes to both centrosomes and ncMTOCs, but is frequently not the sole contributor to microtubule nucleation *in vivo*. For example, γ-TuRC accounts for only a portion of microtubule growth in differentiated skin cells, intestinal cells, and neurons (S. Wang et al., 2015; Sallee et al., 2018; Yau et al., 2014; Sánchez-Huertas et al., 2016). In addition to γ-TuRC, the CAMSAP/Patronin/Nezha family of proteins emerged more recently as a conserved microtubule minus end stabilizer (Goodwin & Vale 2010; Meng et al., 2008; Baines et al., 2009). Unlike γ-TuRC, this family of proteins appears to specifically regulate non-centrosomal microtubules (Akhmanova & Hoogenraad 2015), although the degree to which this protein family is required in microtubule stabilization appears to vary by cell type (S. Wang et al., 2015; Richardson et al., 2014; Yau et al., 2014). Ninein/NOCA-1 is a proposed microtubule anchoring protein, and although it has not been shown to directly bind microtubule minus ends, it likely works in combination with other proteins to anchor microtubules (Mogensen et al., 2000; Lechler & Fuchs 2007; S. Wang et al., 2015). Strikingly, depletion of γ-TuRC, PTRN-1/Patronin, and NOCA-1 simultaneously in *C. elegans* intestinal epithelial cells did not affect the gross morphology or density of non-centrosomal microtubule associated with the apical membrane (Sallee et al., 2018). Thus, essential ncMTOC components have yet to be determined.

Unlike ncMTOCs, essential centrosome components have been defined over the last two decades, a feat made possible by both genetic and proteomic approaches (Bauer et al., 2016; Arslanhan et al., 2020). Neither approach has been used systematically to identify ncMTOC components, perhaps due to a lack of clear phenotypes to screen for and the difficulty in purifying these enigmatic structures. Indeed, studies that probe the protein composition of subcellular compartments traditionally involve biochemical methods such as cell fractionation or immunoprecipitations, which present severe limitations for studying ncMTOCs; biochemical properties specific to ncMTOCs remain largely unknown, making it impossible to fractionate ncMTOCs without high levels of contamination. Additionally, traditional techniques require physical disruption of cells or whole organisms leading to a loss of *in vivo* spatial information of the native cell state and ephemeral interactions between proteins. These issues can be circumvented by biotin ligase-based proximity labeling (PL) techniques, which fuse a protein of interest to a promiscuous labeling enzyme that orchestrates the covalent attachment of biotin to proximal proteins in living cells. Biotinylated proteins can then be isolated and identified by mass spectrometry, providing a “snapshot” of spatial relationships and interaction networks in their native state. PL enzymes historically had slow labeling times (10-18 hours labeling time required in cell culture) and required incubation at 37°C making these techniques impossible to use in a fast-developing organisms grown at cooler temperatures (Roux et al., 2012). Faster acting peroxidase-based labeling (e.g. APEX) also present limitations for intact organisms because of H2O2 toxicity and limited accessibility of the necessary biotin phenol substrate (Reinke et al., 2017). We provided the first demonstration of biotin ligase-based PL in *C. elegans* using the newly-engineered PL enzyme mutant TurboID which presents faster labeling kinetics and retains catalytic activity at lower temperatures than all other existing biotin ligases (Branon et al., 2018). Given all of the advantages this method presents, we used TurboID to identify ncMTOC components in living *C. elegans*.

*C. elegans* embryonic intestinal cells are well-suited for studying ncMTOCs as non-centrosomal microtubules are organized exclusively at the apical membrane starting at embryonic stages and the centrosome is inactivated as an MTOC. The *C. elegans* intestine is derived from a single ‘E’ blastomere that undergoes four rounds of divisions, giving rise to a 16-celled primordium of polarized cells (E16) whose apical membranes all face a common midline (M) that marks the future lumen of the 20-celled (E20) intestine (Figure S1A). During intestinal cells divisions, the centrosome functions as an MTOC, but upon polarization at E16, MTOC function is reassigned to the apical membrane and an apical ncMTOC is established in each cell (Figure S1B) (Feldman & Priess 2012; Yang & Feldman 2015).

To probe the composition of the apical ncMTOC, we expressed TurboID fused to PTRN-1 in embryonic intestinal cells. We identified 69 proteins proximal to PTRN-1, validated the localization of 5 proximal interactors, and focused on identifying functions of two conserved proteins: VAB-10B/MACF1 and WDR-62/H24G06.1, a protein we identify as a homolog of vertebrate WDR62. We report that VAB-10B and WDR-62 specifically localize to the apical ncMTOC in differentiated intestinal cells and in other epithelia but not to the centrosome. Using tissue-specific degradation, we found that these factors regulate non-centrosomal microtubule arrays independently and apparently through distinct mechanisms; loss of VAB-10B resulted in disorganized microtubules and delayed γ-TuRC localization, while loss of WDR-62 decreased microtubule numbers and abolished γ-TuRC localization. VAB-10B and WDR-62 act downstream of cell polarity as the apical polarity protein PAR-3 is properly localized following depletion of VAB-10B or WDR-62, and conversely depletion of PAR-3 mislocalized WDR-62. Finally, we found a role for actin in anchoring microtubules and speculate that this role is downstream of VAB-10B and/or WDR-62. Our data support a model where two essential MTOC functions, microtubule growth and anchoring, are regulated independently by WDR-62 and VAB-10B, respectively. Together, this work identifies components required for ncMTOC formation, highlights that ncMTOCs can be different in composition from centrosomes, and illustrates that ncMTOCs can be composed of different functional modules.

## Results

### Tissue-specific expression of TurboID in *C. elegans* identifies proximity interactors of PTRN-1/Patronin

To identify components of the apical ncMTOC in *C. elegans* embryonic intestinal cells, we sought to identify proteins proximal to PTRN-1, a microtubule minus end binding protein that localizes exclusively to the apical ncMTOC in embryonic intestinal cells and exhibits robust localization throughout development (Figure 1A). We ectopically expressed HA-tagged TurboID fused to PTRN-1 using a promoter that drives expression in intestinal cells (‘PTRN-1::TurboID^gut^’). To control for non-specific biotinylation, we ectopically expressed TurboID alone in parallel using the same promoter (‘TurboID^gut^’). We also analyzed a wild-type (N2) strain not expressing TurboID to control for endogenously biotinylated proteins (Figure 1B).

**Figure 1.**
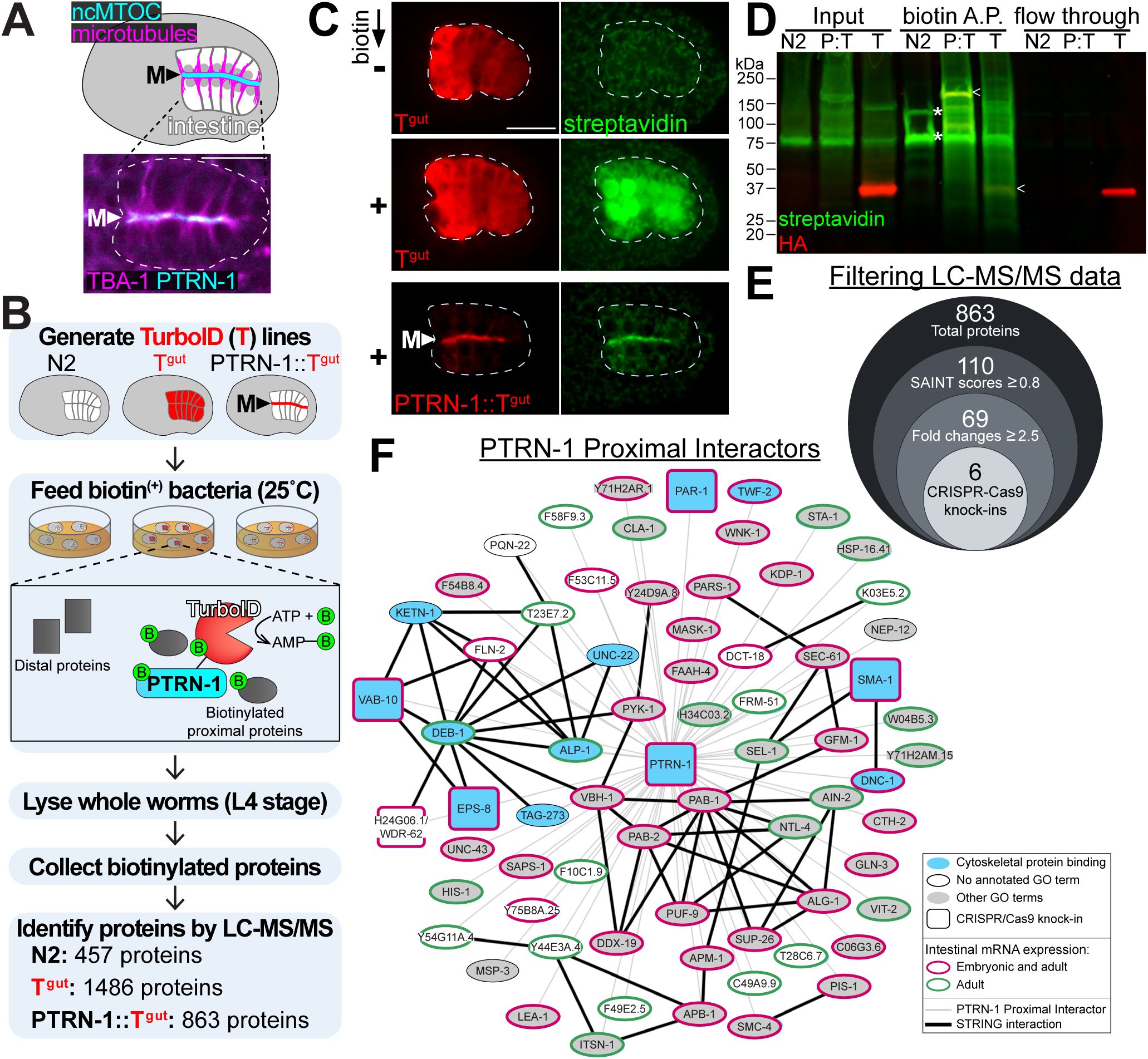
Identification of PTRN-1 proximity interactors. A) Dorsal view of *C. elegans* bean stage embryo indicating the localization of the apical ncMTOC and microtubules in the polarized intestine. Inset shows live imaging of mCherry::TBA-1/α-tubulin and PTRN-1::GFP in intestine (white dotted line). ncMTOC site at the intestinal midline (‘M’) is indicated (arrowhead). B) Experimental workflow for TurboID proximity labeling in the *C. elegans* intestine: wild-type (N2), cytoplasmic TurboID (‘T^gut’^), and TurboID targeted to the ncMTOC (‘PTRN-1::T^gut’^). C) Immunofluorescence imaging in intestine of TurboID (anti-HA, red) and biotinylated proteins (streptavidin-488) for indicated genotypes. Scale bar = 10 μm. D) Immunoblotting for TurboID (anti-HA, red) and biotinylated proteins (streptavidin-488, green) in whole worm lysates (Input) of N2 (wild-type), PTRN-1::TurboID^gut^ (‘P:T’), or TurboID^gut^ (‘T’) that were subjected to pull-downs with streptavidin-covered magnetic beads (biotin A.P.). TurboID (arrowheads) and endogenously biotinylated proteins (asterisks) are marked. Proteins not captured by streptavidin covered beads were also probed (flow through). Representative data from three independent experiments are presented. E) Diagram of workflow to filter LC-MS/MS data and corresponding number of proteins for each step. F) Interaction network of 69 PTRN-1 proximal interactors. The GO term “cytoskeletal protein binding” was significantly enriched (*p* = 5.3e-10). The interaction *p*-value was calculated via Search Tool for the Retrieval of Interacting Genes/Proteins (STRING, *p* = 8.06e-10, black lines); interactions were assessed from co-expression, experimental evidence, databases, and text-mining.

To initially examine biotinylation activity patterns, we fixed and stained embryos with fluorophore-conjugated streptavidin, which binds specifically to biotinylated proteins (Figure 1C). TurboID^gut^-expressing embryos from worms maintained on biotin-producing bacteria exhibited strong biotinylation signal in the intestine, and biotinylation signal was dramatically reduced in embryos from worms maintained on biotin-deficient bacteria, confirming that excess biotin can be delivered through feeding. Embryos expressing PTRN-1::TurboID^gut^ exhibited apically enriched biotinylation activity, indicating that TurboID can mediate regional and spatial control of biotinylation.

Biochemical characterization of protein biotinylation in whole worm lysates by streptavidin blot showed that a wide range of proteins was biotinylated in worms expressing PTRN-1::TurboID^gut^ (Figure 1D) or TurboID^gut^ compared to the endogenously biotinylated proteins found in wild-type worms. Affinity purification of biotinylated proteins using streptavidin-covered beads successfully enriched for biotinylated proteins, including self-biotinylated TurboID^gut^ and PTRN-1::TurboID^gut^. Importantly, biotinylated proteins were detected from whole worm lysates, bypassing the need to perform tissue dissections or fractionation of the ncMTOC structure. Additionally, the lack of biotinylation signal from the flow-through fraction revealed that streptavidin-covered beads captured all biotinylated proteins.

Following identification of biotinylated proteins by mass spectrometry, the 863 total proteins from the PTRN-1::TurboID^gut^ dataset were filtered for SAINT scores (Choi et al., 2011) above or equal to 0.8 and fold changes above or equal to 2.5 when compared to TurboID^gut^ and wild-type (N2) controls (Table S1, Figure 1E). The 69 resulting ‘PTRN-1 proximal interactors’ included VAB-10B (a spectraplakin) and SMA-1 (a β-spectrin), homologs of which have been shown to immunoprecipitate with Patronin/CAMSAP in other systems (Ning et al., 2016; Khanal et al., 2016; King et al., 2014). The isolation of homologs of known PTRN-1 interactors highlights the effectiveness of our approach despite our stringent filtering which could have eliminated lower abundance proteins or proteins with dual cytoplasmic and apical localization. The absence of proteins known to localize to the ncMTOC (e.g. γ-TuRC and NOCA-1) indicates either the restricted labeling radius of PTRN-1::TurboID^gut^, a limitation of TurboID to biotinylate proteins due to a possible shortage of available lysines, a shortage of biotin, or an inability of TurboID to reach proteins that are sterically inaccessible.

We cross-referenced our list of proximal interactors with other datasets (Figure 1F, see Methods). First, we compared our dataset to available RNA seq datasets (Kaletsky et al., 2018; Hashimshony et al., 2015) which showed us that PTRN-1 proximity interactors were highly enriched for genes expressed in the intestine: 1) 39 of the 69 proximity interactors begin mRNA expression in the embryonic intestine and continue into adulthood; and 2) the remaining proximity interactors exhibit mRNA expression in adult intestinal cells with the exception of 6 interactors. These data show that our method enriches TurboID labeling activity in a tissue-specific manner, although with our approach, we cannot pinpoint the developmental stages at which labeling occurred. Additionally, the 69 PTRN-1 proximal interactors were significantly enriched for STRING interactions (*p* = 8.06e-10), suggesting detection of components that exist in interaction or genetic networks. PTRN-1, a protein with microtubule-and actin-associated functions, should in theory be proximal to other proteins with cytoskeleton functions. Consistently, 11 of the 69 genes were statistically enriched for biological function GO term “cytoskeletal protein binding” (*p* = 5.3e-10). Lastly, 57 of the 69 proteins have predicted human orthologs, suggesting that our knowledge of ncMTOC composition is far from complete across organisms.

### Proximity interactors exhibit localization patterns similar to PTRN-1

To further validate that biotinylation activity was occurring at the apical ncMTOC, we probed the localization of 5 PTRN-1 proximal interactors that begin mRNA expression in the embryonic intestine (Hashimshony et al., 2015). Using CRISPR/Cas-9, we inserted a fluorescent tag at the endogenous locus encoding VAB-10B, PAR-1, SMA-1, EPS-8A, and H24G06.1 (Figure S2). These proteins have not been previously shown to interact biochemically or genetically with PTRN-1 in *C. elegans*, but all of their respective homologs have cytoskeleton-related functions. Like PTRN-1, all five proteins localized to the apical membranes of the embryonic intestine (Figure 2A), further underscoring the effectiveness of our proximity labeling approach to identify proximal proteins.

**Figure 2.**
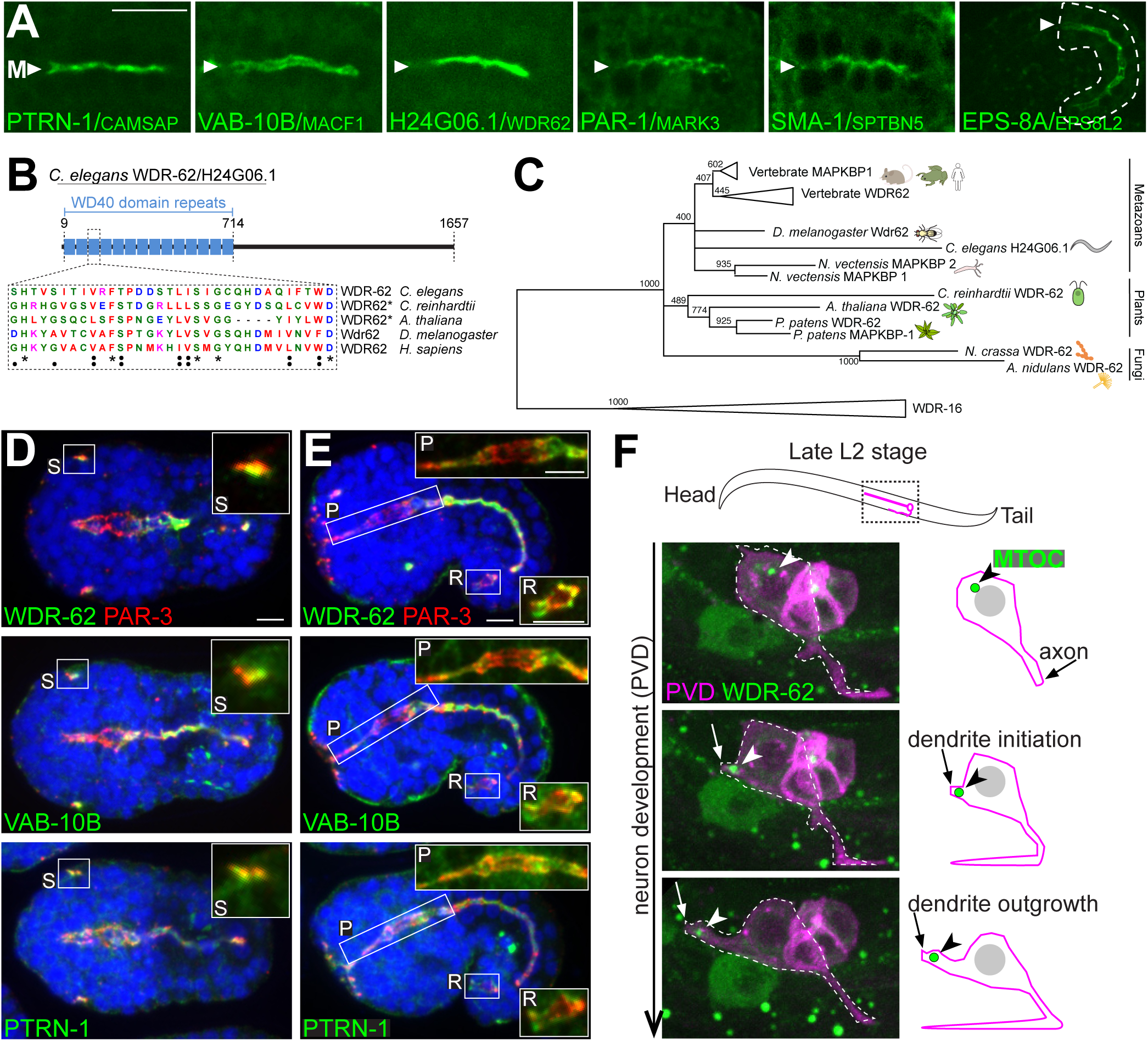
Localization patterns of PTRN-1 proximity interactors. A) Dorsal view live imaging of endogenously ZF::GFP tagged PTRN-1 proximity interactors localizing to the midline (‘M’) of the bean stage polarized intestine. Scale bar = 10 μm. Note EPS-8A is first expressed at a later ‘comma’ stage of intestinal development (see Figure S3). B) Diagram of the domain structure of WDR-62/H24G06.1, indicating the positions of predicted WD40 domain repeats (Ma et al. 2019). Sequence conservation highlighted for single WD40 repeat. C) Collapsed phylogenetic tree from the WD40 repeat region of eukaryotic WDR62 homologs indicates that *C. elegans* H24G06.1 is a WDR62 family member. Maximum likelihood support values are indicated for conserved regions within WD40 repeats. Scale bar represents 0.4 substitutions per site. See also Figure S4. D-E) Immunofluorescence imaging of endogenous PAR-3 (red) and WDR-62::ZF::GFP, VAB-10B::ZF::GFP or PTRN-1::GFP (*α*GFP, green). Insets are higher-magnification views of sensilla (‘S’, ventral embryo view), pharynx (‘P’), and rectum (‘R’). Scale bars = 5 μm. F) Live imaging of WDR-62::ZF::GFP (green) in the developing PVD neuron (magenta). Arrowhead indicates WDR-62 localization to the presumptive ncMTOC (Liang et al. 2020), as represented in adjacent cartoon.

We focused our attention on two proteins: VAB-10B, a spectraplakin, and H24G06.1, an uncharacterized protein. The *vab-10* locus is the only spectraplakin locus in *C. elegans* and encodes two major isoforms, VAB-10A and VAB-10B, that differ in their tissue expression and domain structures; unlike VAB-10B, VAB-10A is not expressed in intestinal cells, and although both VAB-10A and VAB-10B include an N-terminal actin binding domain, they differ in a C-terminal intermediate filament and microtubule binding domain, respectively (Figure S2) (Bosher et al., 2003; Gally et al., 2016). Using amino-acid sequence homology and phylogenetic analyses, we identified H24G06.1 (hereafter called WDR-62) as a divergent homolog of the primary autosomal recessive microcephaly (MCPH) protein WDR62 and its paralog MAPKBP1 (Pervaiz & Abbasi 2016) (Figure 2B, C, S4, Table S2). Mutations in *WDR62* are the second most common cause of MCPH in human patients (Morris-Rosendahl et al., 2015), but the cellular roles of WDR62 remain incompletely understood with the vast majority of studies focusing on a role for WDR62 at centrosomes (Yu et al., 2010; Bogoyevitch et al., 2012; Chen et al., 2014; Ramdas Nair et al., 2016; Jayaraman et al., 2016; Sgourdou et al., 2017). By investigating the endogenous localization of WDR-62 tagged at either the N-or C-terminus, we first detected *wdr-62* expression in the embryonic intestine, but saw no evidence of WDR-62 centrosomal localization, including in dividing intestinal precursor cells (Fig. S2B). Additionally, we found that WDR-62 does not appear to be maternally expressed and is not found in any of the rapidly dividing cells of the early embryo (data not shown). Moreover, we found that, like in embryonic intestinal cells, WDR-62 localizes to the apical surfaces of other epithelial cell types as indicated by colocalization with the conserved apical polarity protein PAR-3/PAR3 (Figure 2D, E). Epithelial expression and apical localization were similar for WDR-62, VAB-10B, and PTRN-1, for example at the apical dendrite tip of developing amphid sensilla neurons (Figure 2D, ‘S’) and in pharyngeal and rectal cells (Figure 2E, ‘P’, ‘R’). As human and mouse WDR62 are known to play prominent roles in neuronal development (Shohayeb et al., 2018), we investigated the localization of WDR-62 in the developing PVD neuron, a highly branched nocioceptor. Intriguingly, we found that WDR-62 localizes to the tip of the outgrowing PVD dendrite (arrowhead, Figure 2F), a site that was recently shown to contain a migrating ncMTOC enriched for microtubules and γ-TuRC (Figure 2F) (Liang et al., 2020). Our discovery of WDR-62 in many polarized cell types provided a unique opportunity to study its role beyond the centrosome.

### VAB-10B and WDR-62 are required to form non-centrosomal microtubule arrays

The localization patterns of endogenous VAB-10B and WDR-62 and their proximity to PTRN-1 prompted us to investigate non-centrosomal microtubule-related functions for these proteins. To do this, we employed the ZF/ZIF-1 tissue-specific protein degradation system whereby proteins containing a ZF degron tag are targeted for degradation by the E3 ligase adapter ZIF-1 (Armenti et al., 2014; Sallee et al., 2018). We inserted a ZF::GFP tag into the endogenous locus of *vab-10b* and *wdr-62* using CRISPR/Cas9 (Figure S2). The resulting “ZF-tagged” proteins were degraded by ectopically expressing ZIF-1 under the control of the *elt-2* promoter, which drives ZIF-1 expression in the intestine starting at the E2-E4 stage several hours before ncMTOC formation and allows for intestine-specific depletion of these targets (“VAB-10B^gut(-)^,” “WDR-62^gut(-)^”, Figure S2).

In control embryos, mCherry::TBA-1/α-tubulin was strongly enriched at apical membranes (“M,” Figure 3), signifying apical microtubule organization as has been previously described (Figure 3A, B) (Feldman & Priess 2012). By contrast, mCherry::TBA-1 was significantly reduced at apical membranes in VAB-10B^gut(-)^ and WDR-62^gut(-)^ embryos compared to control (Figure 3A, B). Residual microtubules in WDR-62^gut(-)^ embryos could be due to other microtubule regulators or unannotated WDR-62 isoforms that exclude the C terminal ZF degron (Figure S2B). Co-depleting VAB-10B and WDR-62 resulted in an even more severe depletion of mCherry::TBA-1 at the apical membranes compared to control (Figure 3A,B) or to VAB-10B^gut(-)^ and WDR-62^gut(-)^ single depletion embryos, suggesting that VAB-10B and WDR-62 organize non-centrosomal microtubules in separate pathways.

**Figure 3.**
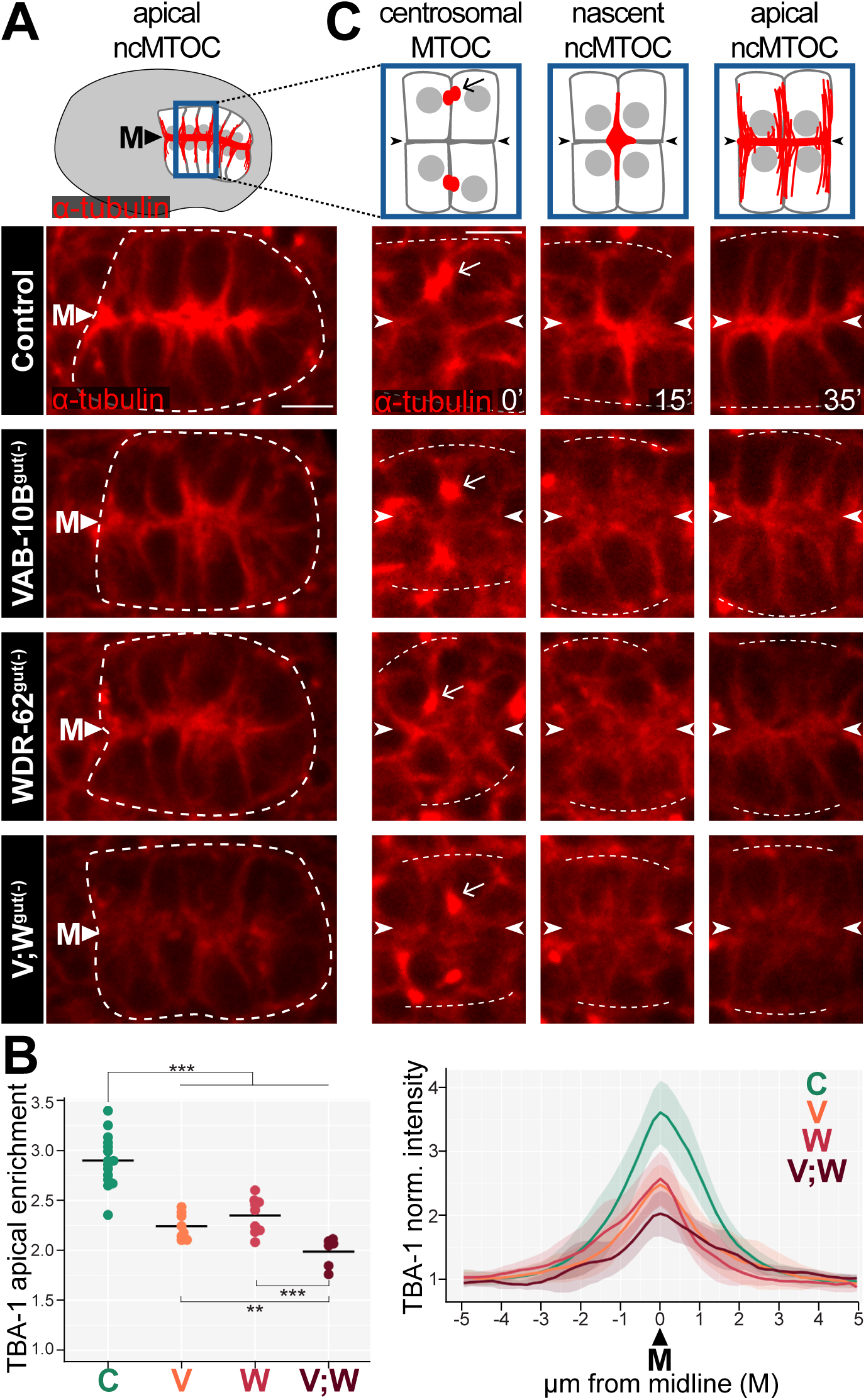
VAB-10B and WDR-62 are required for non-centrosomal microtubule organization. A) Dorsal view of intestine (white dotted line) from live imaging of mCherry::TBA-1/*α*-tubulin in control embryos or embryos with intestine-specific depletion of VAB-10B (VAB-10B^gut(-)^), WDR-62 (WDR-62^gut(-)^), or simultaneous depletion of VAB-10B and WDR-62 (V;W^gut(-)^). Cartoon of embryonic intestine (top) represents microtubules (red), nuclei (gray), midline (‘M’) in control embryos. B) Left: Quantification of mCherry::TBA-1 fluorescent signal enrichment at the apical midline where horizontal black lines show mean; \*\**p* = 0.0073, \*\*\**p* < 0.00099. Right: Normalized fluorescent signal along a line perpendicular to the intestinal midline; lighter shading indicates standard deviation. For both graphs, Control (‘C’) n = 15; VAB-10B^gut(-)^ (‘V’) n = 8; WDR-62^gut(-)^ (‘W’) n = 9; V;W^gut(-)^ (‘V;W’) n = 8. C) Cartoons depict dorsal view of four intestinal cells during ncMTOC establishment in control embryos. Live imaging of mCherry:TBA-1 in cells exiting E8-E16 division (t = 0’), polarizing E16 cells (t = 15’), and newly polarized E16 cells (t = 35’). Arrow points to presumptive centrosomal TBA-1 signal near the lateral membrane of adjacent non-sister cells. Arrowheads point to future apical midline. Scale bars = 5 μm.

In order to determine the time at which microtubule phenotypes arise, we tracked microtubule localization during ncMTOC formation, a process that occurs ∼30 minutes after the E8-E16 division. After embryonic intestinal cells exit mitosis at E16, the centrosome is inactivated as an MTOC and a nascent ncMTOC localizing γ-TuRC and microtubules forms de novo that will move to and spread across the midline of the intestine to establish the apical ncMTOC (Feldman & Priess 2012). We have previously found that microtubules follow a stereotyped pattern of movement, shown here by live-imaging of mCherry:TBA-1 in control embryos (Figure 3C) (Feldman & Priess 2012). Centrosomal microtubules (arrow) from the E8-E16 division localize to lateral membranes (t = 0’). As the centrosome is inactivated as an MTOC, non-centrosomal microtubules emerge near lateral membranes and move toward the presumptive apical surfaces, first localizing at cell vertices (t =15’) before finally spreading along the midline (‘M’, arrowheads) and orienting along the apicobasal axis (t = 35’). In VAB-10B^gut(-)^, WDR-62^gut(-)^, and V;W^gut(-)^ embryos, centrosomal microtubules (Figure 3C, arrow, t=0’) appeared unaffected, but non-centrosomal microtubules were diffuse during their initial formation (t = 15’) and reduced at the apical membrane, with the greatest reduction seen in V;W^gut(-)^ embryos (t = 35’). These data indicate that VAB-10B and WDR-62 are required to organize non-centrosomal microtubules beginning at ncMTOC formation.

### WDR-62 and VAB-10B control dynamic microtubule growth and localization

To provide deeper insight into the microtubule patterning roles we found for VAB-10B and WDR-62, we investigated microtubule dynamics in VAB-10B^gut(-)^ and WDR-62^gut(-)^ embryos. Analysis of endogenous localization patterns of the microtubule plus end binding protein EBP-2/EB3 showed overall numbers of dynamic microtubules and their growth speed and direction. In control embryos, EBP-2::GFP comets originated at the apical membrane of each cell and moved toward the basal surface (Video S1). Images of a single time point and a 10-second time projection illustrate a high enrichment of comets at the intestinal midline and their ‘fountain-like’ pattern of movement away from the apical surface, respectively (Figure 4A). From the single time point images, it became evident that removal of VAB-10B or WDR-62 resulted in decreased enrichment of EBP-2::GFP at the midline (Figure 4A). Furthermore, the time projection images showed significant differences in the number of comet trajectories in WDR-62^gut(-)^ intestines as compared to VAB-10B^gut(-)^, representing another key departure of their respective phenotypes.

**Figure 4.**
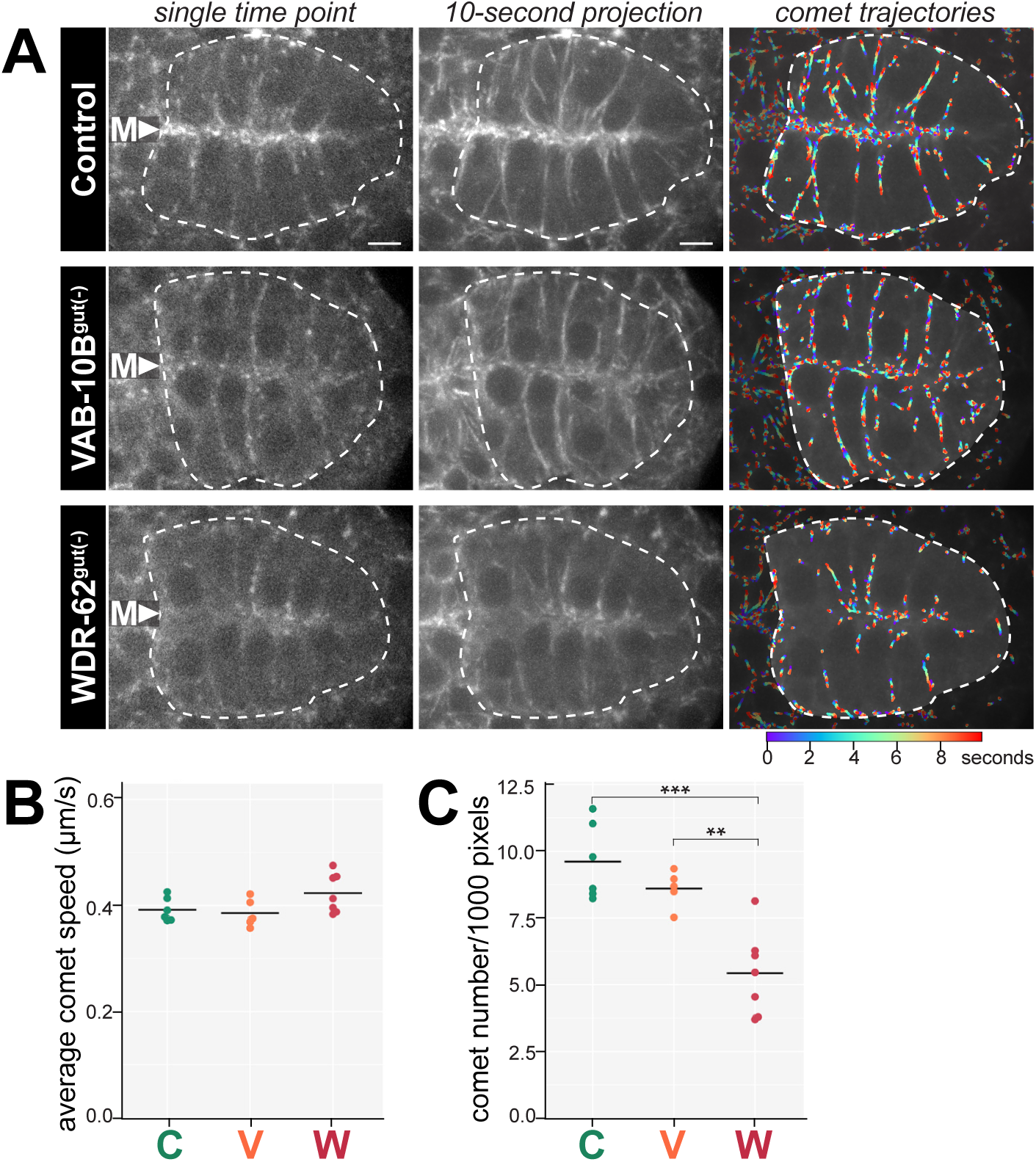
Depletion of VAB-10B or WDR-62 differentially affect dynamic microtubules. A) Dorsal view of live bean-stage polarized intestines (white dotted lines) from rapid time-lapse imaging of EBP-2/EB1::GFP in indicated genotypes (see Video S1) represented as a single time point, 10-second time projection, or overlaid trajectories as indicated. The color map indicates time frame for comet trajectory lines; blue is 0 seconds and red is the last time point. Midline (‘M’) is indicated. Scale bars = 5 μm. B, C) Computationally tracked comet trajectories were quantified to measure average comet speed (B) and average comet number per 1000 pixels in the intestine (C): control (‘C’) n = 7; VAB-10B^gut(-)^ (‘V’) n = 5; WDR-62^gut(-)^ (‘W’) n = 7; \*\**p* = 0.0012; \*\*\**p* = 0.0004.

We quantified our observations of EBP-2 comets in different genotypes using custom-built segmentation and tracking algorithms (Figure 4B, see Methods). Overall, the average comet speed did not differ between genotypes (Figure 4B). However, we observed significant differences in the total number of comets between genotypes. Control and VAB-10B^gut(-)^ embryos had a significantly higher number of intestinal EBP-2 comets compared to WDR-62^gut(-)^ embryos and control embryos did not statistically differ from VAB-10B^gut(-)^ (Figure 4C). These data indicate that unlike VAB-10B depletion, loss of WDR-62 reduced the number of dynamic microtubules emanating from the apical surfaces. Although VAB-10B depletions did not change the number of dynamic microtubules, the EBP-2 signal was less enriched at the apical surfaces compared to control and comet trajectories appeared more disorganized. In all, these data suggest that VAB-10B does not function in microtubule growth, but rather to anchor microtubules to the apical membrane, whereas WDR-62 plays a major role in the production of microtubules.

### VAB-10B and WDR-62 recruit microtubule minus end proteins to the apical membrane

Given the roles we found for VAB-10B and WDR-62 in microtubule organization and dynamics, we sought to understand their roles in localizing other MTOC components, including microtubule minus end-binding proteins γ-TuRC and PTRN-1. In polarized intestinal cells, the γ-TuRC component GIP-1/GCP3 localized to both the apical ncMTOC (arrowhead, Figure 5A) and to inactive centrosomes (joined double arrows, Figure 5A, (Feldman & Priess 2012)). The centrosomal localization of GIP-1 was unaffected in VAB-10B^gut(-)^ and WDR-62^gut(-)^ embryos (joined double arrows, Figure 5A), suggesting that inactive centrosomes are not affected by these protein depletions. In contrast, removal of VAB-10B resulted in mislocalized non-centrosomal GIP-1 (arrowheads, Figure 5A). Even more strikingly, WDR-62^gut(-)^ embryos exhibited a drastic decrease of non-centrosomal GIP-1 signal (arrowheads, Figure 5A).

**Figure 5.**
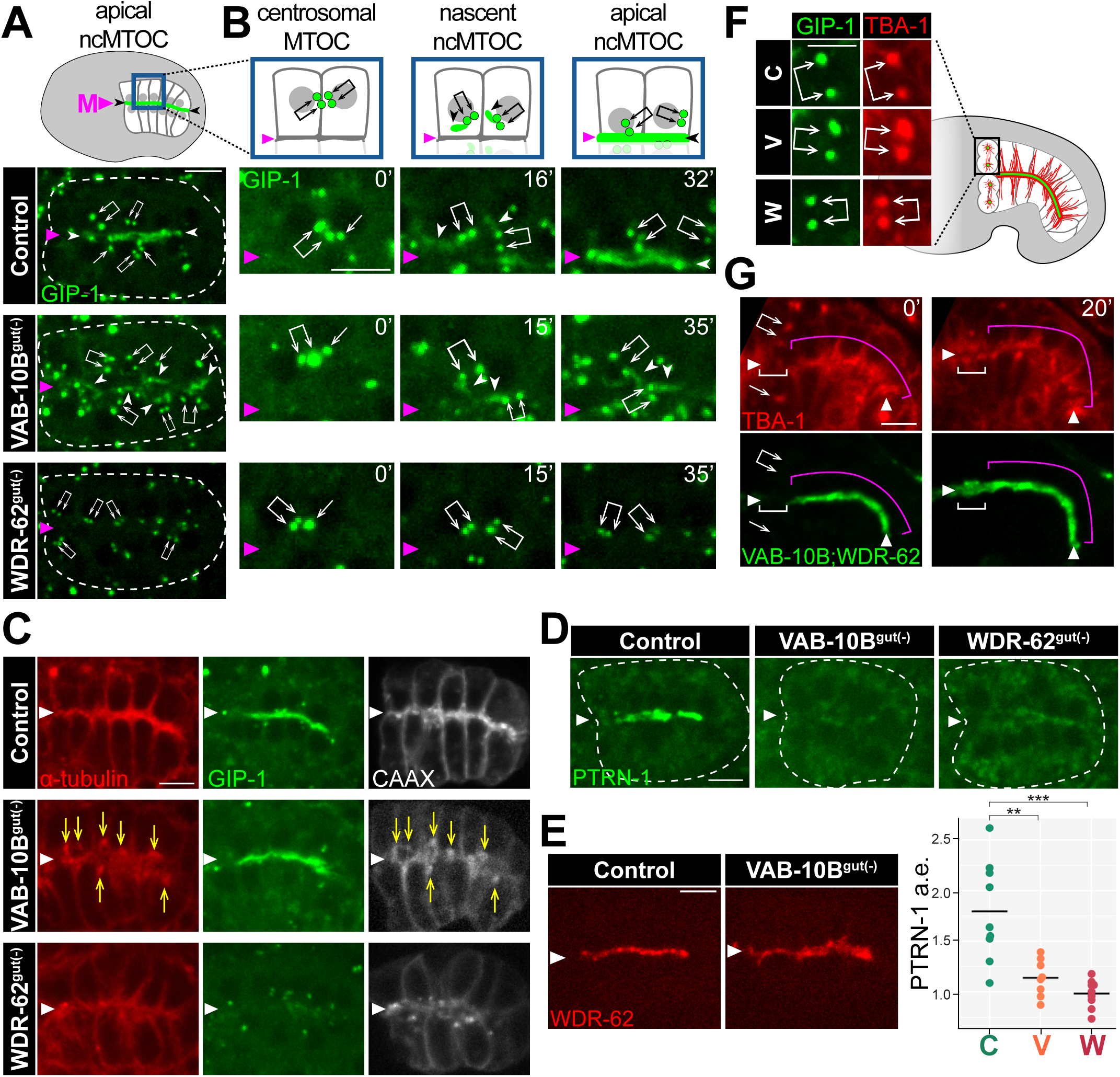
VAB-10B and WDR-62 differentially recruit microtubule minus end proteins to the ncMTOC. A, B) Dorsal view live imaging of endogenous GFP::GIP-1/GCP3 localized to centrosomes (single arrow or joined white arrows) and as a non-centrosomal pool (white arrowheads) in embryonic intestines in control, VAB-10B depletion (VAB-10B^gut(-)^), or WDR-62 depletion (WDR-62^gut(-)^). Midline is marked with a magenta triangle (‘M’). Cartoon represents GFP::GIP-1 (green) and nuclei (gray) of control (see Figure S1). B) Magnified view of two intestinal cells exiting the E8-E16 division (t = 0’), during polarization (t = 15’, t = 16’), and just after apical polarization (t = 32’ or t = 35’). Time lapse n = 3. C) Live imaging of mCherry::TBA-1/*α*-tubulin, GFP::GIP-1, and membrane marker BFP::CAAX. Yellow arrows indicate position of mislocalized TBA-1 puncta. D) Live imaging of endogenous PTRN-1::GFP localization in polarized intestines with quantification of apical enrichment (a.e., bottom) for control (‘C’) n = 9, VAB-10B^gut(-)^ (‘V’) n = 8, and WDR-62^gut(-)^ (‘W’) n = 9; \*\*\**p*= 0.0009; \*\**p*= 0.0038. E) Live imaging of WDR-62::RFP localization after VAB-10B depletion. F) mCherry::TBA-1 and GFP::GIP-1 localization at centrosomes (joined white arrows) in ‘comma stage’ intestines when anterior cells re-enter the cell cycle. F) Live imaging of mCherry::TBA-1 and both endogenous VAB-10B::ZF::GFP and WDR-62::ZF::GFP in comma stage intestines undergoing anterior cell divisions (t = 0’). Following division, anterior cells re-polarize (t=20’). Centrosomes (joined double arrows or single arrow), midline (white triangles) and differentiated cells (magenta bracket) are indicated. Scale bars = 5 μm.

To better understand the precise time at which these phenotypes arise, we performed time-lapse imaging of GIP-1 during ncMTOC formation. In control embryos, GIP-1 localized to centrosomes at lateral membranes between adjacent non-sister E16 cells as they exit mitosis (joined double arrows, t = 0’, Figure 5B, Figure S1, (Feldman & Priess 2012)). Next, ‘plumes’ of non-centrosomal GIP-1 appeared adjacent to centrosomes, (arrowheads, t = 16’) which subsequently coalesced at the midline (t = 32’). In VAB-10B^gut(-)^ embryos, centrosomal GIP-1 localization appeared normal at mitotic exit (t = 0’) and throughout polarization; however, although non-centrosomal GIP-1 plumes were still apparent (t = 15), they showed a severe delay in their coalescence at the midline, suggesting that VAB-10B links GIP-1 to the apical ncMTOC. In WDR-62^gut(-)^ embryos, the centrosomal pool of GIP-1 appeared normal (t = 0’), but non-centrosomal GIP-1 plumes neither formed (t = 15’) nor appeared at the midline (t = 35’), suggesting that WDR-62 plays a role different from that of VAB-10B, possibly in initially forming the non-centrosomal microtubules that can then recruit γ-TuRC.

At a later developmental stage, around 1 hour after polarization, apically localized GIP-1 and TBA-1 remained reduced in WDR-62^gut(-)^ embryos (Figure 5C). In contrast to WDR-62^gut(-)^ embryos and to earlier phenotypes, VAB-10B^gut(-)^ embryos showed normal apical GIP-1 localization at this stage, but reduced and disorganized microtubules as compared to control (Figure 5C). Indeed, we frequently (n = 8/10) observed aberrant TBA-1 accumulations (arrows) near the apical membrane. These foci were not coincident with centrosomes as indicated by GIP-1 localization but instead colocalized with intestine-specific BFP targeted to membranes by the Ras CAAX motif (‘BFP:CAAX’, VAB-10B^gut(-)^, Figure 5C), which has been shown to localize to various membrane compartments including the plasma membrane and vesicles of the secretory pathway (Choy et al., 1999). These data highlight that γ-TuRC is differentially regulated by WDR-62 and VAB-10B, and furthermore that γ-TuRC is not always a faithful marker of microtubule organization.

In contrast to the non-centrosomal γ-TuRC localization defects we observed in WDR-62^gut(-)^ or VAB-10B^gut(-)^ embryos, we found that apical PTRN-1 signal was drastically decreased upon depletion of either VAB-10B or WDR-62 (Figure 5D). This result is consistent with several reports of spectraplakins targeting Patronin/CAMSAP to ncMTOCs in various cell types (Ning et al., 2016; Nashchekin et al., 2016; Zheng et al., 2020) and is the first report of a role for WDR62 family proteins in PTRN-1 localization.

The different microtubule and γ-TuRC localization phenotypes following VAB-10B and WDR-62 depletion and the more severe microtubule phenotype in the double mutant as compared to either single mutant suggested that these two proteins function in parallel pathways. Thus, we predicted that VAB-10B and WDR-62 would independently localize to the apical membrane. Indeed, WDR-62 was properly localized to the apical surface following VAB-10B depletion (Figure 5G). These data further support the hypothesis that WDR-62 and VAB-10B act in parallel pathways to regulate microtubule and minus end protein organization at the apical ncMTOC.

Finally, we questioned whether the microtubule abnormalities caused by WDR-62 and VAB-10B depletions were specific to the apical ncMTOC. After intestinal cell polarization, four intestinal cells re-enter mitosis and microtubules and γ-TuRC are removed from the apical ncMTOC and instead localize to active centrosomes (Figure 5E, (Yang & Feldman 2015)).

Although WDR-62 and VAB-10B depletions perturbed the accumulation of microtubules and GIP-1 signal at the apical ncMTOC, centrosomal accumulation was not affected (Figure 5E), suggesting that the microtubule defects caused by loss of WDR-62 and VAB-10B are specific to the apical ncMTOC. Consistent with this result, both VAB-10B and WDR-62 leave the apical membrane during mitosis (white bracket, Figure 5F) but do not accumulate at the centrosomes (arrows), while the localization of these proteins in adjacent non-dividing intestinal cells did not change (pink bracket, t = 0’). Following mitosis, VAB-10B and WDR-62 returned to the apical membrane (white bracket, t = 20’), suggesting that VAB-10B and WDR-62 track with and function at non-centrosomal rather than centrosomal microtubules.

### The conserved PAR polarity complex localizes with and organizes PTRN-1, WDR-62, and VAB-10B

The movement of the nascent ncMTOC to the midline during E16 intestinal cell polarization is coincident with and controlled by the conserved apical polarity protein PAR-3 (Figure S1B, (Feldman & Priess 2012)). Non-centrosomal microtubules and plumes of γ-TuRC form near and move to the midline with polarity proteins destined for the apical surface, and depletion of PAR-3 inhibits the apical localization of γ-TuRC and microtubules (Feldman & Priess 2012; Achilleos et al., 2010). To determine if other ncMTOC components exhibit similar localization patterns, we investigated the localization of PTRN-1, WDR-62 and VAB-10B relative to PAR-3 during polarization. PTRN-1, WDR-62, and VAB-10B initially appeared as foci on lateral membranes with PAR-3, moved together with PAR-3 toward the midline (Figure 6A, A’), and then spread across the midline to subsequently increase in intensity after cells established an apical surface (Figure 6A’’). Importantly, none of these proteins colocalized with centrosomes and instead appeared as part of a separate ncMTOC structure as it formed de novo (Figure S1C).

**Figure 6.**
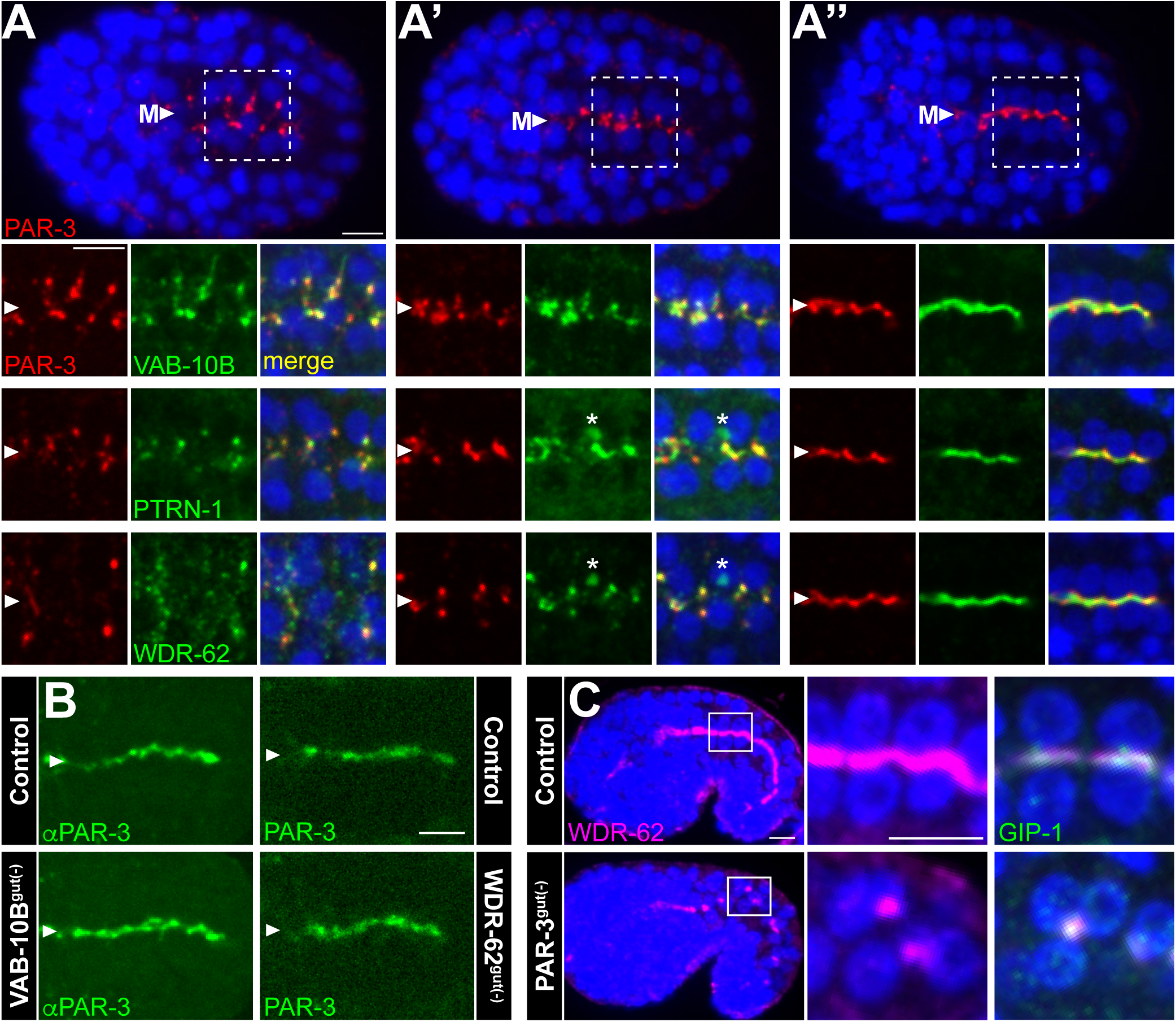
VAB-10B and WDR-62 interface with apical polarity determinants. (A-A’’) Dorsal views from immunofluorescence imaging of endogenous PAR-3 (*α*PAR-3, red) or GFP from WDR-62::ZF::GFP, VAB-10B::ZF::GFP, or PTRN-1::GFP (*α*GFP, green). Higher-magnification views of white boxed region from intestinal cells are shown. Arrowhead marks the intestinal midline (‘M’). Asterisks mark intruding green signal from ventral germ cells. B) Immunofluorescence (*α*PAR-3, left) or live imaging (PAR-3:mCherry, right) of endogenous PAR-3 in indicated genotypes. C) Immunofluorescence imaging of endogenous WDR-62::RFP (magenta, αRFP), γ-TuRC (*α*GIP-1, green), and nuclei (blue, DAPI), in control or PAR-3^gut(-)^ comma stage embryos. Scale bars = 5 μm.

The observation that apical polarity factor PAR-3 colocalizes with VAB-10B and WDR-62 during intestinal polarization led us to ask whether polarity cues control VAB-10B and WDR-62 localization or vice versa. PAR-3 localization was unaffected in VAB-10B^gut(-)^ or WDR-62^gut(-)^ embryos (Figure 6B), suggesting that neither VAB-10B nor WDR-62 control the most upstream aspects of apical polarization. In contrast, depletion of PAR-3 specifically in intestinal cells (PAR-3^gut(-)^) mislocalized WDR-62 into large puncta away from the midline that co-localized with GIP-1 (Figure 6C), indicating that PAR-3 directs WDR-62 to the apical surface and tracks with the ncMTOC upon its mislocalization.

### Actin filaments regulate non-centrosomal microtubules

Spectraplakins are thought to function as linkers between actin and non-centrosomal microtubules by recruiting microtubule-bound Patronin/CAMSAP to the actin cortex of epithelial cells (Ning et al., 2016; Nashchekin et al., 2016). The spectraplakin VAB-10B has both an actin and microtubule binding domain, and we initially hypothesized that VAB-10B organizes non-centrosomal microtubules by tethering them to the actin cortex of intestinal cells. We confirmed the apical enrichment of actin filaments with a transgene encoding YFP::ACT-5, an actin gene that is highly expressed in the intestine (Figure 7A, t=0’, (MacQueen et al., 2005)). We then perturbed the actin network by culturing embryos in the actin polymerization inhibitors Latrunculin A (LatA) and Cytochalasin D (CytoD). *C. elegans* embryos are normally surrounded by an impermeable eggshell and vitelline membrane that can be permeabilized on demand using a laser, allowing the introduction of inhibitors with temporal precision. LatA-treated embryos showed a robust depletion of YFP::ACT-5 from apical surfaces, illustrating the effectiveness of the drug treatment in inhibiting actin polymerization in the intestine (Figure 7A, t = 20’). We treated embryos expressing endogenously-tagged WDR-62, VAB-10B, or PTRN-1 with actin inhibitors (Figure 7B-D). In the presence of actin inhibitors, apical surfaces became gnarled, consistent with previous reports that actin polymerization defects cause apical membrane constrictions, again indicating the effectiveness of our inhibitor treatments (van Furden et al., 2004). Surprisingly and contrary to our initial hypothesis, neither WDR-62, VAB-10B, nor PTRN-1 were removed from the apical surface following treatment with actin inhibitors. Conversely, we were unable to observe any changes in apical actin localization in VAB-10B^gut(-)^ or WDR-62 ^gut(-)^ embryos (Figure 7E). We next treated embryos expressing endogenously tagged GFP::TBB-2/*β*-tubulin (Figure 7F, Video S2) or a transgene encoding mCherry::TBA-1/*α*-tubulin (n = 8, data not shown) with actin inhibitors when microtubules were enriched at the apical surface (t = 0’). Strikingly, in the presence of actin inhibitors, microtubules appeared to be released from apical surfaces throughout the intestine as indicated by the gradual removal of TBB-2 (t = 10’-25’). The anterior intestinal cells that undergo an additional round of mitosis following polarization (t = 25’, asterisk) were still able to recruit microtubules to the centrosome to build the mitotic spindle suggesting a specific non-centrosomal role for actin in microtubule anchoring.

**Figure 7.**
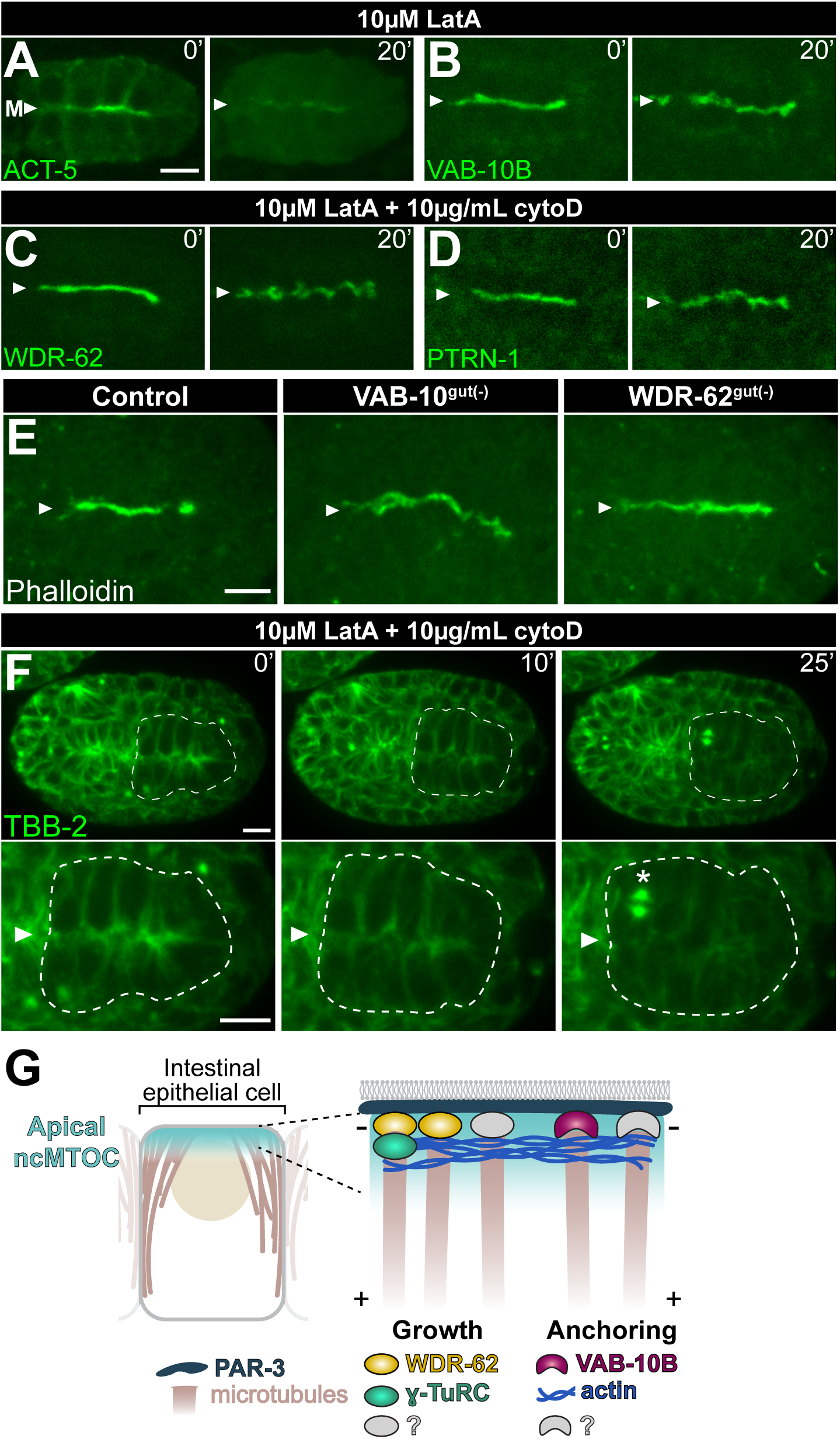
Actin filaments regulate non-centrosomal microtubules. (A-D) Time-lapse imaging of dorsal view in bean stage embryos expressing YFP::ACT-5 (n = 3), or endogenously tagged VAB-10B::ZF::GFP (n = 4), WDR-62:ZF::GFP (n = 3), PTRN-1::GFP (n = 3) beginning several seconds after eggshell permeabilization (t = 0’) in the presence of the indicated actin inhibitor for 20 minutes (t = 20’). E) Dorsal view immunofluorescence staining of actin filaments (phalloidin-488) in indicated genotypes. (F) Time-lapse imaging of GFP::TBB-2/β-tubulin with intestine (white dotted line), midline (‘M’), and mitotic spindle (asterisk) indicated; n = 3; see Video S2. G) Model of apical ncMTOC composition and function. Scale bars = 5 μm.

## Discussion

Using biotin-based proximity labeling coupled with a tissue-specific degradation system in *C. elegans* differentiated intestinal cells, we addressed fundamental knowledge gaps in ncMTOC biology. This study presents technical and conceptual advances: 1) TurboID is an effective method for identifying proximity interactors in a living multicellular organism, in this case proximal interactors at an ncMTOC; 2) Spectraplakin VAB-10B and WD40 repeat protein WDR-62 are two essential ncMTOC components with distinct functions; and 3) ncMTOCs can be composed of functionally distinct modules that separate microtubule growth and anchoring (Figure 7G).

We showed the effectiveness of biotin-based proximity labeling in *C. elegans*, which allowed us to present the first ncMTOC proteome in polarized cells. While we limited this study to the functions of VAB-10B and WDR-62, we identified additional PTRN-1 proximity interactors with diverse functional GO terms, including ‘actin binding’ and ‘RNA binding’. Given the central role we found for actin in anchoring non-centrosomal microtubules in the embryonic intestine, it will be necessary to further investigate the role of proximity interactors that also interact with actin. RNA binding proteins have not previously been shown to localize to ncMTOCs, however the existence and function of RNA at the centrosome has a long history and so this class of proteins is an interesting category for further investigation (Marshall & Rosenbaum 2000; Alliegro et al., 2006; Sepulveda et al., 2018).

Our analysis of PTRN-1 proximity interactors revealed previously uncharacterized proteins, including H24G06.1, a homolog of vertebrate WDR62 and its paralog MAPKBP1. The cellular roles for MAPKBP1 remain poorly understood, although a localization study showed MAPKBP1 at mitotic spindle poles (Macia et al., 2017). Previous studies of WDR62 have largely focused on centrosomal roles in mitosis as it localizes at spindle poles in dividing cultured cells and is required for the stability of spindle microtubules and for timely cell cycle progression (Yu et al., 2010; Chen et al., 2014). WDR62 also has centrosomal functions in interphase cells as depletion of Wdr62 in mouse embryonic fibroblasts leads to centriole duplication defects, and the fly ortholog Wdr62 maintains interphase centrosomal MTOC activity in neuroblasts, potentially through the stabilization of centrosomal microtubules (Jayaraman et al., 2016; Ramdas Nair et al., 2016). By using differentiated *C. elegans* intestinal cells as a model, we found that WDR-62 has additional non-mitotic, non-centrosomal roles as an essential component of the apical ncMTOC. These roles are likely not unique to *C. elegans* as *Wdr62* depletion disrupted the apical complex proteins in mouse cortical neurons through an unexplored mechanism (Jayaraman et al., 2016). Furthermore, our phylogenetic analyses suggest that the ancient function of WDR62 proteins may be rooted in more basic microtubule functions as WDR62 is conserved in organisms that do not have centrosomes such as *A. thaliana* and *P. patens*.

Our tissue-specific depletion studies provide evidence for separate functional modules controlling growth and anchoring within ncMTOCs. We found that WRD-62 functions in a microtubule growth as WDR-62 depletion significantly reduced the number of dynamic microtubules emanating from the apical ncMTOC. Conceptually, microtubule growth could be directed by proteins whose function impacts microtubule stabilization and/or nucleation, however distinguishing these functions *in vivo* can be difficult. Mutational analysis shows that WDR62 binds microtubules through its WD40 domains providing further evidence for its direct role in stabilizing and/or nucleating microtubules (Bogoyevitch et al., 2012; Lim et al., 2015). Wdr62 depletion in Drosophila neuroblasts reduced microtubule localization to interphase centrosomes and microtubules were more sensitive to microtubule depolymerizing treatments; these data are again consistent with Wdr62 acting as either a stabilizer or nucleator (Ramdas Nair et al., 2016). Given that WDR-62 depletion abolished localization of the conserved microtubule nucleator γ-TuRC, we propose that WDR-62 functions in microtubule nucleation. As the microtubule defects previously reported from γ-TuRC depletion in intestinal cells (Sallee et al., 2018) were minimal compared to the defects we report from WDR-62 depletion, the role of WDR-62 in nucleation would be to directly nucleate γ-TuRC-independent microtubules or to indirectly promote nucleation through the recruitment of currently undiscovered nucleation factors.

The spectraplakin VAB-10B appears to function in a second independent module which imparts microtubule anchoring. Although spectraplakins are well known for binding the MT plus-end binding protein EB1 and the microtubule lattice (Sun et al., 2001; Kodama et al., 2003; Honnappa et al., 2009), the Drosophila spectraplakin Shot, and one of the two human spectraplakins ACF7, have been shown to interact with the microtubule minus end through Patronin and CAMSAP3, respectively (Nashchekin et al., 2016; Ning et al., 2016).

Spectraplakins are largely thought to crosslink non-centrosomal microtubules to the cortex by interacting with Patronin/CAMSAP3 at the minus end of non-centrosomal microtubules and anchoring them to cortical actin filaments. Consistent with a role for spectraplakins in microtubule anchoring, we found that VAB-10B localizes near microtubule minus ends and its depletion does not change the number of dynamic microtubules, but rather their localization. We also found that VAB-10B recruits PTRN-1, as in other systems, however the microtubule anchoring function of VAB-10B is not dependent on PTRN-1, as depletion of PTRN-1 had no effect on microtubules in the embryonic intestine (Sallee et al., 2018). Furthermore, although CAMSAP proteins have domains that promote binding to microtubule minus ends, we found that loss of apical microtubules through actin inhibition had no effect on PTRN-1 localization. We therefore propose that at the apical ncMTOC, VAB-10B recruits PTRN-1 independently of microtubules and VAB-10B anchors microtubules independently of PTRN-1. Exactly how VAB-10 anchors microtubules, either directly through its known MT binding domains or as a platform for other microtubule regulators is currently unknown.

An understanding of how VAB-10B and WDR-62 becomes apically localized will reveal mechanistic aspects of ncMTOC establishment, as we found these proteins are essential components of the apical ncMTOC. Given previous reports, we expected actin to serve as a platform for VAB-10B localization to the apical membrane (Applewhite et al., 2010; Ning et al., 2016; Suozzi et al., 2012). Indeed, VAB-10B depletion caused apical surfaces to become gnarled similar to the apical constriction phenotypes previously observed after actin perturbation (Figure 7E, Figure 5G) suggesting that VAB-10B and actin function in the same pathway (van Furden et al., 2004). However, neither VAB-10B nor WDR-62 localization were affected following actin inhibition. In contrast and to our surprise, the same pharmacological inhibition of the actin network liberated microtubules from the apical surface. This strong displacement of microtubules following inhibition of actin polymerization suggests that actin is required downstream of VAB-10B and/or WDR-62. While removing VAB-10B or WDR-62 did not change actin localization, we speculate that these proteins could localize a small population of actin that is unresolvable against the vast apically enriched pools of actin that will ultimately contribute to the terminal web. Alternatively, or in addition, VAB-10B and WDR-62 could work together to localize actin, consistent with the fact that VAB-10B has an actin binding domain and knockdown of WDR62 in mouse oocytes disrupted the formation of the actin cap (Y.S. Wang et al., 2020). Although we do not yet know the nature of these potential interactions, actin was recently shown to directly bind to γ-TuRC and perturbing this interaction inhibited γ-TuRC nucleation activity (Liu et al., 2020), suggesting a direct role for actin in microtubule growth.

The cellular origin of non-centrosomal microtubules and by extension the function of ncMTOCs is a topic of debate and likely varies by cell type. Different non-mutually exclusive models have been proposed that differ in their involvement of the centrosome. For example, microtubules grown at the centrosome could be released and captured at a non-centrosomal site. As an extension of this model, nucleators from the centrosome could be transferred to the ncMTOC. Alternatively, ncMTOCs could be nucleating and anchoring microtubules without involvement of centrosome intermediates. Our data suggest that ncMTOCs predominantly nucleate and anchor microtubules using molecules that do not have centrosomal origins. First, we showed that the composition of the apical ncMTOC differs from the centrosome, as neither VAB-10B nor WDR-62 localize to centrosomes. Second, depleting VAB-10B and WDR-62 specifically affected non-centrosomal γ-TuRC and microtubules, but did not perturb the localization of γ-TuRC or microtubules to the centrosome. These results further underscore the model that centrosomes and ncMTOCs contain different essential components and argue against a centrosomal origin for non-centrosomal microtubules and their regulators. As we have previously found a role for the centrosome and microtubules in apical ncMTOC establishment (Feldman & Priess 2012), we favor a model where the centrosome, perhaps through its astral microtubules, creates an environment that promotes the timely formation of the nascent ncMTOC.

Our data lead to a model where apical microtubules are grown and anchored by two centrosome-independent modules, separating these two microtubule functions at ncMTOCs (Figure 7G). This type of division of labor has been suggested for other MTOCs (Nashchekin et al., 2016; S. Wang et al., 2015; Muroyama et al., 2016). For example, centrosomes in differentiating keratinocytes have separate γ-TuRC complexes thought to nucleate (CDK5RAP2-γ-TuRC) or anchor (Nedd-γ-TuRC) microtubules (Muroyama et al., 2016). Instead of involving the same protein in different functional subcomplexes as at the centrosome, we suggest that WDR-62 regulates microtubule growth and VAB-10B regulates microtubule anchoring. A small portion of microtubules are still able to grow when WDR-62 is depleted, suggesting that other WDR-62 isoforms are not targeted by our degron system or opening up the exciting possibility that additional microtubule nucleators exist.

How an ncMTOC is built de novo at the onset of differentiation and how it forms a non-membrane bound functional unit is an open question. We found in cells depleted of the apical polarity regulator PAR-3 that WDR-62 and GIP-1 still coalesce yet form into a large mislocalized patch within each intestinal cell. Thus, in PAR-3 depleted cells, the ncMTOC can still form but at the wrong subcellular site. What causes ncMTOC molecules to coalesce? The ncMTOC could be a phase-separated structure, as PTRN-1 and several PTRN-1 proximity interactors contain intrinsically disordered regions. Alternatively, and consistent with the colocalization of displaced microtubules with membrane puncta in VAB-10^gut(-)^ embryos, membranous structures could serve as a platform for ncMTOC assembly as Golgi, Golgi outposts, and endosomes have all been shown to associate with non-centrosomal microtubules (Ori-McKenney et al., 2012; Efimov et al., 2007; Hehnly & Doxsey 2014; Liang et al., 2020). Thus PAR-3 and other polarity regulators could merely position an otherwise autonomous structure.

Diverse non-centrosomal microtubule arrays have long been observed in a wide range of differentiated cell types. Our proximity labeling approach coupled with a tissue specific degradation method has uncovered key ncMTOC components in polarized epithelial cells that will serve as the foundation for our understanding of the fundamental mechanisms cells use to generate specific microtubule patterns.

## Acknowledgements

We thank Ashby Morrison, Kristy Red-Horse, Asako Sugimoto, Karen Oegema, Dan Dickinson, and Bob Goldstein for providing reagents, strains, and for CRISPR advice and protocols. We also thank members of the Feldman lab and Andrew McKay for helpful discussions about the manuscript. Some of the nematode strains used in this work were provided by the *Caenorhabditis* Genetic Center, which is funded by the NIH Office of Research Infrastructure Programs (P40 OD010440). Mass Spectrometry was performed at Stanford University (Vincent Coates Foundation Mass Spectrometry Laboratory, Stanford University Mass Spectrometry). This work was supported by an NIH New Innovator Award DP2GM119136-01 and R01GM133950 awarded to J.L.F. A.D.S. was supported by NIH Training Grant 2T32GM007276. T.C.B. was supported by Dow Graduate Research and Lester Wolfe Fellowships. C.J.W. and K.S. are investigators of the Howard Hughes Medical Institute. A.P. was supported by Rubicon Science 2018-1 research program (019.182EN.018) partly financed by the Dutch Research Council (NWO). K. S. was supported by NIH NS082208. A.Y.T. was supported by NIH R01 DK121409.

## Author contributions

Conceptualization, J.L.F. and A.D.S.; Methodology, A.D.S., J.L.F., T.B., A.Y.T., A.P., C.J.W.; Formal Analysis, A.D.S.; Investigation, A.D.S., J.L.F., L.E.C., M.A.P., T.B., X.L., A.P.; Writing - Original Draft, A.D.S. and J.L.F.; Writing – Review & Editing, A.D.S., J.L.F., L.E.C., M.A.P., T.B., A.Y.T., C.J.W.; Visualization, J.L.F., A.D.S., A.P., L.E.C., and X.L.; Supervision, J.L.F., K.S., A.Y.T., C.J.W.; Funding Acquisition, J.L.F., A.D.S., A.Y.T., A.P., C.J.W., K.S.

## STAR methods

### C. elegans strains and maintenance

Nematodes were cultured and manipulated as previously described (Brenner 1974). Unless otherwise noted, strains were cultured and maintained at 20°C on E. coli OP50 bacteria. To deplete animals of excess biotin, animals were grown on washed biotin auxotrophic E. coli (MG1655bioB:kan) (Ortega-Cuellar et al., 2010). The strains used in this study are listed in Table S3.

Extrachromosomal arrays used in this study:

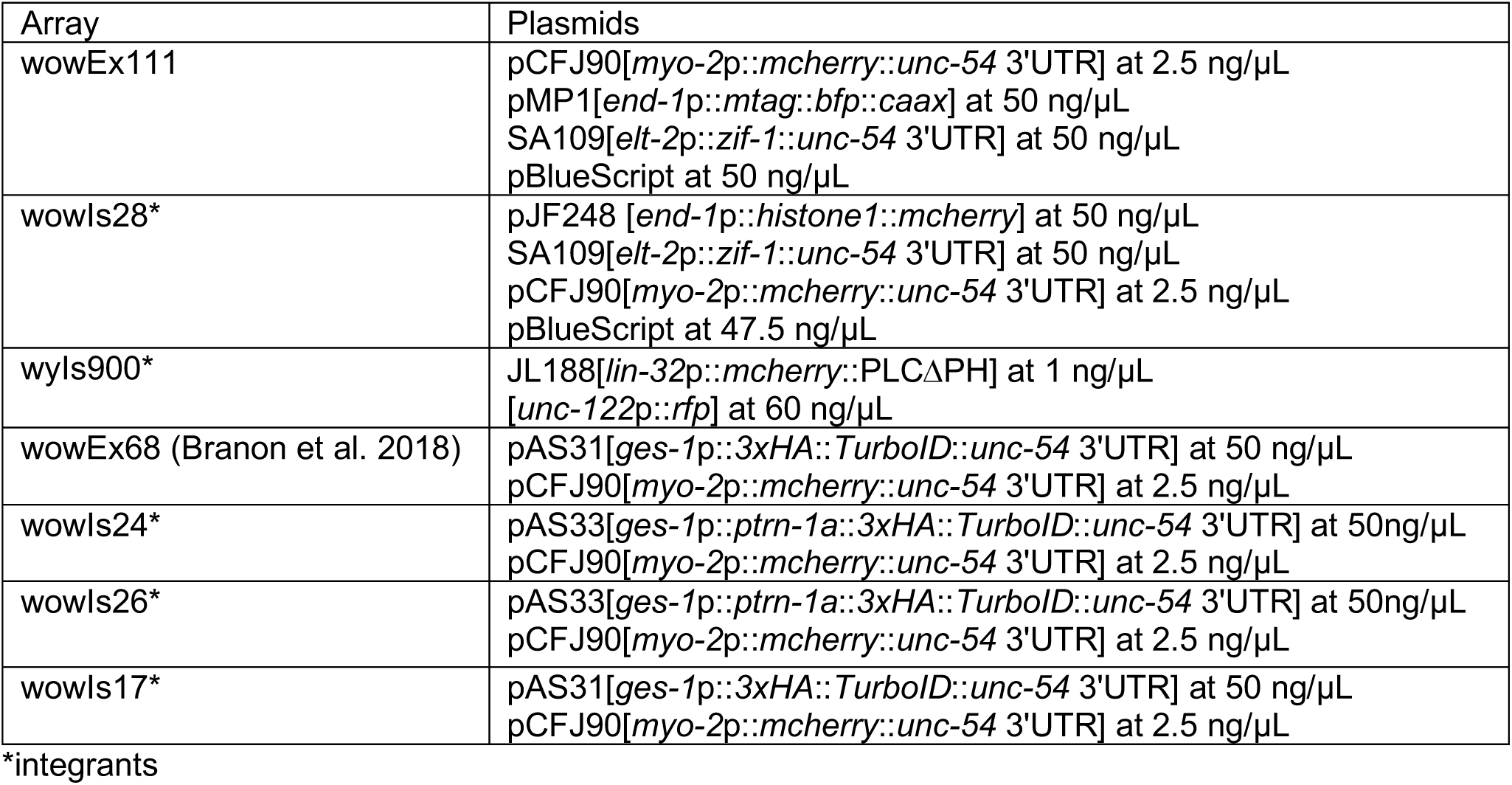

| Array | Plasmids |
| --- | --- |
| wowEx111 | pCFJ90[myo-2p::mcherry::unc-54 3'UTR] at 2.5 ng/μL<br>pMP1[end-1p::mtag::bfp::caax] at 50 ng/μL<br>SA109[elt-2p::zif-1::unc-54 3'UTR] at 50 ng/μL<br>pBlueScript at 50 ng/μL |
| wowls28* | pJF248 [end-1p::histone1::mcherry] at 50 ng/μL<br>SA109[elt-2p::zif-1::unc-54 3'UTR] at 50 ng/μL<br>pCFJ90[myo-2p::mcherry::unc-54 3'UTR] at 2.5 ng/μL<br>pBlueScript at 47.5 ng/μL |
| wyls900* | JL188[lin-32p::mcherry::PLCΔPH] at 1 ng/μL<br>[unc-122p::rfp] at 60 ng/μL |
| wowEx68 (Branon et al. 2018) | pAS31[ges-1p::3xHA::TurboID::unc-54 3'UTR] at 50 ng/μL<br>pCFJ90[myo-2p::mcherry::unc-54 3'UTR] at 2.5 ng/μL |
| wowls24* | pAS33[ges-1p::ptrn-1a::3xHA::TurboID::unc-54 3'UTR] at 50ng/μL<br>pCFJ90[myo-2p::mcherry::unc-54 3'UTR] at 2.5 ng/μL |
| wowls26* | pAS33[ges-1p::ptrn-1a::3xHA::TurboID::unc-54 3'UTR] at 50ng/μL<br>pCFJ90[myo-2p::mcherry::unc-54 3'UTR] at 2.5 ng/μL |
| wowls17* | pAS31[ges-1p::3xHA::TurboID::unc-54 3'UTR] at 50 ng/μL<br>pCFJ90[myo-2p::mcherry::unc-54 3'UTR] at 2.5 ng/μL |

\*integrants

### TurboID Proximity-Dependent Protein Labeling Method

#### Plasmids and strains

*C. elegans* codon-optimized ligase gene TurboID (containing 3 worm introns present in GFP) is as previously published (Branon et al., 2018). All expression constructs were cloned into pJF241 to produce plasmids pAS31 (*ges-1p::3xha::turboID::unc-54*) and pAS33 (*ges-1p::ptrn-1a::3xha::turboID::unc-54*). Transgenic worms were generated by injecting 50ng/uL plasmid and 2.5ng/uL of the co-injection marker *myo-2*p::mCherry into day 1 N2 hermaphrodites. Extrachromosomal arrays were integrated with gamma irradiation (total dose of 3800 radians) using a cesium irradiator.

#### Mass spectrometry sample preparation

Embryos of each genotype were plated on XL peptone-rich plates coated in NA22 bacteria and grown until L4 stage at 25°C. Worms were harvested via M9 washes and frozen in 500μL pellets at −80°C and unless otherwise noted, all following steps were done at 4°C. On the day of sample preparation, frozen worm pellets were thawed to 4°C in 600 μL of high SDS RIPA buffer (50 mM Tris-HCl pH 8.0; 150 mM NaCl; 1% SDS; 0.5% sodium deoxycholate; 1% TritonX-100; 1mM PMSF; 1x HALT; 2.5 mg/mL Leupeptin; 5 mg/mL Pepstatin), transferred to bead beating tubes (lysing matrix C, 1.0mm silica spheres, MP Biomedicals), and agitated at 6.5m/s, 20 sec pulse (3x) with 5 minute breaks on 4°C ice (FastPrep-24 Classic). Samples were transferred to sonication tubes and sonicated at 18% at 5 second intervals with 5 second breaks for 50 seconds total in a 4°C bath (Branson Sonifier SFX250). SDS-free RIPA buffer was added to achieve a final SDS concentration of 0.2%. Rotate lysate for 1 hour at 4°C. Samples were transferred to ultracentrifuge tubes (13×51MM), and spun for 30 minutes at 51,000 RPM in a TLA-100.3 ultracentrifuge rotor (Beckman). The liquid layer was collected, avoiding the lipid layer, and 12mg of total input protein was rotated overnight at 4°C with washed streptavidin beads (Pierce Streptavidin Magnetic Beads, Thermo Fisher Scientific, #88817) at a ratio of 8μg protein/1μL beads. After binding, the samples were washed for 2 min. each with the following solutions and indicated number of repeats: RIPA buffer (2x), 1M KCl (1x), 0.1M Na2CO3 (1x), 2M urea (1x), 4M urea (1x), RIPA (2x), PBS (5x).

#### Western Blotting

Lysate was loaded onto a 4-20% Mini-PROTEAN TGX PAGE gel (Bio-Rad), transferred to a nitrocellulose membrane (0.4 μm, Bio-Rad). Blots were blocked in 5% milk PBST solution, probed with anti-HA (1:5000, rat monoclonal, Roche), and detected with secondary antibody (1:5000, goat anti-rat IRDye 680RD, Licor) and streptavidin-IRDye (1:5000, 800CW, Licor). Blots were imaged on LI-COR Odyssey CLx.

#### Mass Spectrometry

Biotinylated samples were processed for mass spectrometry on streptavidin beads. Beads were re-suspended in 50mM ammonium bicarbonate and then reduced with 10 mM DTT at 55°C for 5 min., followed by 25 min. at room temperature. Alkylation was performed with 30mM acrylamide for 30 min. at room temperature. Digestion was performed with Trypsin/LysC (Promega) in the presence of 0.02% protease max (Promega) in a standard overnight digest at room temperature on a head-over-head mixer. The digestion reaction was quenched with 1% formic acid and peptides were de-salted with C18 Monospin reversed phase columns (GL Sciences). De-salted peptides were dried in a speed vac. Samples were reconstituted in 20μl reconstitution buffer (2% acetonitrile with 0.1% Formic acid), 2μl of which was injected on the instrument.

Mass spectrometry experiments were performed using an Orbitrap Fusion Tribrid mass spectrometer (Thermo Scientific, San Jose, CA) with liquid chromatography using a Nanoacquity UPLC (Waters Corporation, Milford, MA). For a typical LCMS experiment, a flow rate of 450 nL/min was used, where mobile phase A was 0.2% formic acid in water and mobile phase B was 0.2% formic acid in acetonitrile. Analytical columns were prepared in-house with an I.D. of 100 microns pulled to a nanospray emitter using a P2000 laser puller (Sutter Instrument, Novato, CA). The column was packed using C18 reprosil Pur 1.8 micron stationary phase (Dr. Maisch) to a length of ∼25 cm. Peptides were directly injected onto the analytical column using a gradient (2-45% B, followed by a high-B wash) of 80min. The mass spectrometer was operated in a data dependent fashion using CID fragmentation for MS/MS spectra generation.

#### Mass spectrometry analysis

MS/MS data were processed using Byonic (Protein Metrics, San Carlos, CA) to identify peptides and infer proteins using the *C. elegans* database from Uniprot. Proteolysis with Trypsin/LysC was assumed to be semi-specific allowing for N-ragged cleavage with up to two missed cleavage sites. Precursor mass accuracies were held within 12 ppm, and 0.4 Da for MS/MS fragments. Proteins were held to a false discovery rate of 1%, using standard approaches. Subsequent analyses were done using Uniprot Protein ID and corresponding spectral count after contaminants were removed. Any Uniprot Protein ID not detected for a given sample was assigned a spectral count value of 0 for the purpose of the analysis. For pairwise comparisons, protein length and spectral counts for each sample were uploaded to crapome.org (Mellacheruvu, Nat Methods 2013, database version v1.1) and the following parameters were used: workflow 3, fold change score options: FC-A(user, default); FC-B(all, stringent), C is control, Saint Expressed SAINT options: Saint Express with user control [(SE)C(user)]; incorporate known data (IKD)=none; number of replicates per bait (NR)=all. SAINT analysis score (Choi et al., 2011) and fold change value for each Uniprot Protein ID were calculated. PTRN-1 proximity interactors were defined by a minimum SAINT score of ≥0.8 and a minimum fold change over each control of ≥2.5. GOTermFinder was used to find shared GO terms with *p*-values indicated in Table S1 (Boyle et al. 2004).

### Dataset queries

Adult RNAseq information was acquired from: http://worm.princeton.edu/ (Kaletsky et al., 2018). GO term analyses were done using GO term finder https://go.princeton.edu/cgi-bin/GOTermFinder (Boyle et al., 2004). To find predicted human orthologs of *C. elegans* genes, OrthoList (Shaye & Greenwald 2011) and Biomart (www.biomart.org) were used.

### CRISPR/Cas9

All CRISPR/Cas9-based insertions were achieved using the self-excising cassette (SEC) method (Dickinson et al., 2015). Plasmid pDD286 was used for insertion of 3xMyc and TagRFP-T or plasmid JF250 was used for insertion of a ZF domain, 3xFLAG, and GFP (Sallee et al., 2018). Cas9 and sgRNAs for each edit were expressed from plasmid pDD162, into which the appropriate sgRNA sequence was added with a Q5 Site-Directed Mutagenesis Kit. For each edit, young adult *zif-1(gk117)* hermaphrodites were injected with the SEC plasmid and Cas9/sgRNA plasmid. Worms were recovered and screened for the expected edit according to published protocols (Dickinson et al., 2015). Sequences for sgRNA and homology arms are listed.

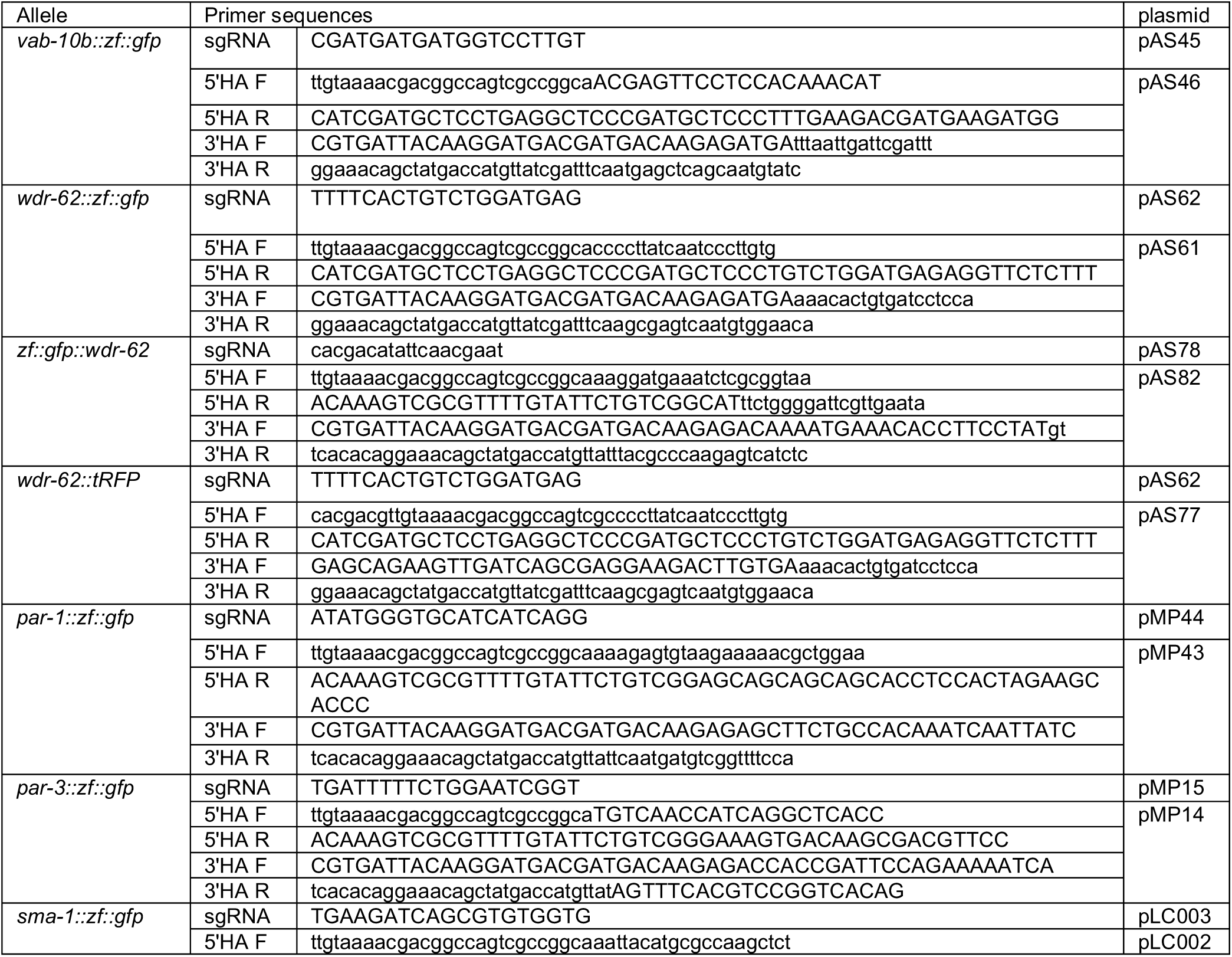
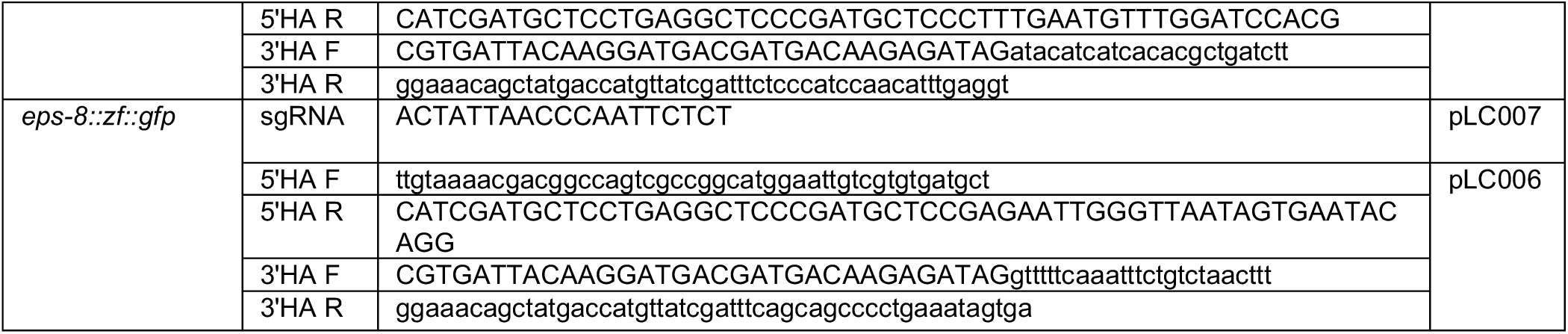

### ZF/ZIF-1 degradation

The ZF/ZIF-1 system was used as previously described (Sallee et al. 2018). As described above (See CRISPR/Cas9), the ZF degron and adjoining GFP were inserted into the endogenous locus of *zif-1(gk117)* hermaphrodites. Exogenous ZIF-1 was expressed from either an extrachromosomal array (wowEx111) or by single-copy insertion as indicated in strain Table S3 (S. Wang et al., 2017). The extrachromosomal array was generated as indicated in extrachromosomal injection table (above). To keep degradation methodology consistent, we used the same array across all strains by mating. Loss of GFP signal is demonstrated in Figure S2.

### Inhibition of actin polymerization

Actin inhibitor experiments were performed as previously described (Feldman & Priess 2012). Briefly, embryos were coated with 0.1% trypan blue and attached to coverslips coated with poly-lysine in Shelton and Bowerman buffer (SGM) containing either 10μM Latrunculin A or a mixture of 10μM Latruculin A with 10ug/mL Cytochlasin D. The coverslip was inverted over a slide with Teflon spacers and 15-20 22.5 μm glass beads to prevent the coverslip from bending under the pressure from the objective (Whitehouse scientific). The eggshell and vitelline membrane of E16-stage embryos were permeabilized using a Micropoint pulse-nitrogen dye laser (Coumarin dye, Andor). Embryos were imaged before and then immediately after laser permeabilization at 5 minute intervals.

### Immunohistochemistry

One-day-old adults were incubated in 1xM9 solution for 4-5 hours and their embryos were removed, fixed, and stained using previously described staining methods (Leung et al., 1999). Embryos were attached to microscope slides coated with poly-lysine and containing Teflon spacers. Slides were frozen on dry ice, embryos were permeabilized by freeze-crack method and submerged in 100% MeOH for 5-10 minutes at −20°C. Embryos were submerged for 5 minutes twice in PBS, then once in PBT (PBS plus 0.1% Tween). Embryos were incubated in primary antibody overnight at 4°C. Embryos were then washed in PBT for 5 minutes three times and then incubated in secondary antibody for 1 hour at 37°C. Embryos were washed once in PBT then twice in PBS, mounted in Vectashield (Vector Laboratories), and stored at 4°C. The following primary antibodies were used: anti-HA primary antibody (Abcam, 1:200), anti-Myc (Abcam, 1:100), anti-GFP (Abcam, 1:200), anti-PAR-3 (DSHB), anti-GIP-1 (GenScript, (Feldman & Priess 2012)). The following secondary antibodies were used: CY3-anti-mouse secondary antibody (Jackson Immunoresearch Laboratories, 1:200), Streptavidin Alexa Fluor 488 (Invitrogen, 1:200), DAPI (Sigma, 1:10,000), 647-anti-rabbit (Jackson Immunoresearch Laboratories, 1:50), 488-anti-mouse (Jackson Immunoresearch Laboratories, 1:200), 488 anti-rabbit (Jackson Immunoresearch Laboratories, 1:200), Cy5 anti-mouse (Jackson Immunoresearch Laboratories, 1:50).

For phalloidin staining, embryos were treated as previously described (van Furden et al. 2004). Briefly, after embryos were permeabilized by freeze-crack method, they were submerged in a solution of 75% methanol and 4% paraformaldehyde for 30 minutes at −20deg. Embryos were washed twice with PBT, 10 minutes each, and incubated at room temperature for 1 hour with Alexa Fluor 488 phalloidin (Invitrogen, 1:25) in a moist chamber, then washed twice for 10 minutes each in PBS, mounted in Vectashield (Vector Laboratories), and stored at 4°C.

### Microscopy

For live imaging, embryos were isolated from young hermaphrodites incubated for 4-5 hours in 1x M9 solution at 20°C. Embryos were mounted on a pad made of 3% agarose dissolved in 1x M9 solution and imaged on a Nikon Ti-E inverted microscope (Nikon Instruments, Melville, NY) with a confocal spinning disk head using a 60x or 100x PLAN APO oil objective (NA = 1.4 or NA = 1.45, respectively) controlled using NIS Elements software (Nikon). Images were captured using an Andor Ixon Ultra back thinned EM-CCD camera controlled using NIS Elements software (Nikon) at a z-sampling rate of 0.5 μm and using 488 nm, 561 nm, 405 nm, or 640 nm imaging lasers. Fixed images were acquired using a 60x PLAN APO oil objective (NA = 1.4) on the above Nikon confocal system. All images were processed in NIS Elements software, Fiji/ImageJ version 2.0, and/or Adobe InDesign.

### Image quantification

#### Analysis considerations

Embryos were carefully chosen based on the appropriate stage of development and with their dorsal side oriented toward the coverslip. The same microscopy imaging parameters were used across genotypes for a given experiment.

#### Quantification of apical fluorescence enrichment

Analysis was done as previously described (Sallee et al., 2018). Bean-stage embryos were chosen for analysis. Using Fiji, a membrane marker, nuclei positions, and distance from primordial germ cells were used to select the plane in the Z axis in which to define the intestinal midline. For each embryo, a sum Z-projection of three slices flanking the midline plane was used for analysis. Boxes 2 μm in width were drawn by hand in intestinal cytoplasmic regions to calculate background fluorescence and one box was drawn at the apical midline to define the region of interest (ROI). For each embryo, apical enrichment was calculated by dividing the mean intensity within the apical ROI by the mean cytoplasmic intensity. To plot line profiles, a 1-μm wide line was hand-drawn perpendicular to the apical midline. Plots for apical enrichment and profile plots were generated in R using ggbeeswarm and ggplot2, respectively. Each profile was normalized by dividing each pixel intensity value by the average intensity 2.5 μm to 5 μm from the midline. The mean profile was calculated with the intensity value for each distance from the midline.

### EBP-2 comet analysis

For the segmentation and tracking of EBP-2 comets, as well as the estimation of the speed and angle of microtubule polymerization, we used a set of custom-built filters and functions in Python.

#### Segmentation of the embryo and the embryonic intestine

For the segmentation of the embryo, an adaptive filter was used on the smoothed image. An intestine-specific BFP::CAAX image was first smoothed using the “gaussian_filter” function from the scipy.ndimage package. The threshold_local filter from the skimage.filters package was then applied on the smoothed image to generate a dynamic threshold. This dynamic threshold, estimated on the basis of the local pixel neighborhood, was used to mask the embryo pixels (pixels higher than the local threshold). The edges of the binary mask were bridged and the holes were filled using the binary_closing and the binary_fill_holes functions from the scipy.ndimage package. An area threshold in pixels was used to remove small masked regions (noise around the embryo). If there were more than one embryo in the field of view, the largest embryo was selected.

#### Segmentation of EBP-2 comets

For the detection and segmentation of the EBP-2 comets, a Laplacian of Gaussian (LoG) filter and adaptive thresholding were applied on the EBP-2::GFP images. The LoG filter was constructed by smoothing the image (scipy.ndimage.gaussian_filter) and then applying a Laplace operator (skimage.filters.laplace). A percentile of the brightest pixels was set as a hard threshold on the LoG image to generate a binary mask. A second binary mask was generated by adaptively thresholding (skimage.filters.threshold_local) the original fluorescence image. The two binary masks were multiplied. As a result, only those pixels that are selected in both masks will correspond to the pixels of the segmented comets. The masked spots in the image were labeled. A minimum area threshold was used to remove small labels that did not correspond to comets. To better separate clustered comets (i.e. a mask label containing more than one comet), we zoomed into mask labels larger than a specified area threshold where we applied a new LoG filter and hard threshold with stricter parameters. The centroid coordinates of the mask labels, each corresponding to one EBP-2 comet, were used as the location of the comet. The segmentation algorithm was applied to all fluorescence images in the stream acquisition.

#### Tracking of the EBP-2 comets

The tracking of the microtubule comets was performed using the distance between comet labels in subsequent frames. A search radius is specified by the user (in pixels) and the algorithm links EBP-2 comet to other comets within the search radius in the next time-point. If more than one comet is detected, the most proximal one is selected. If no comet is detected within the search radius in the next time-point, the algorithm searches for comets in subsequent time-points within 8 frames. This allows for complete comet trajectories even when a few comet positions are missed.

#### Comet trajectory speed estimation

Because the comets are tracked every 100msec, the displacement between adjacent time-points in the stream acquisition can be smaller than the comet localization error. In order to improve the comet displacement statistics, we applied a time-lag of 4 frames in the displacement estimation. As a result, the displacement of the comet position between each frame and 4 frames ahead was calculated. All spurious trajectories that included less than 5 frames were removed from the analysis. The EBP-2 comet extension speed was estimated on the basis of these 4 frame intervals. The same time-lag was applied for estimating the angle of the comet displacements.

### Statistical analysis

For all statistical procedures, R was used to perform one-way ANOVA followed by unpaired Student’s t-tests. The Dunn’s method was applied as a post-hoc test to assess significance. Figure legends indicate *p* values.

### Phylogenetic analysis

Representative WDR-16 and WDR-62 homologs from invertebrates (*Caenorhabditis elegans* H24G06.1a), vertebrates (*Homo sapiens* WDR-16, WDR-62, and MAPKBP-1), and chytrid fungi (*Batrachochytrium dendrobatidis* BDEG_26806 and BATDEDRAFT_35991) were used to identify reciprocal best blast hits in species represented in the phylogenetic tree (Table S2). Number of WD40 repeats was determined by WDSP (Ma et al., 2019). Sequences were aligned using Clustal Omega before trimming to 230 amino acid positions within the WD40 repeat region using Gblocks v0.91 with parameters -b3=15 -b4=2 -b5=a. PhyML 3.0 (Guindon et al., 2010; Lefort et al., 2017) with substitution model (LG+G) selected by Akaike Information Criterion was used to generate support values using 1000 bootstraps.

## Supplemental Figure legends

**Figure S1.**
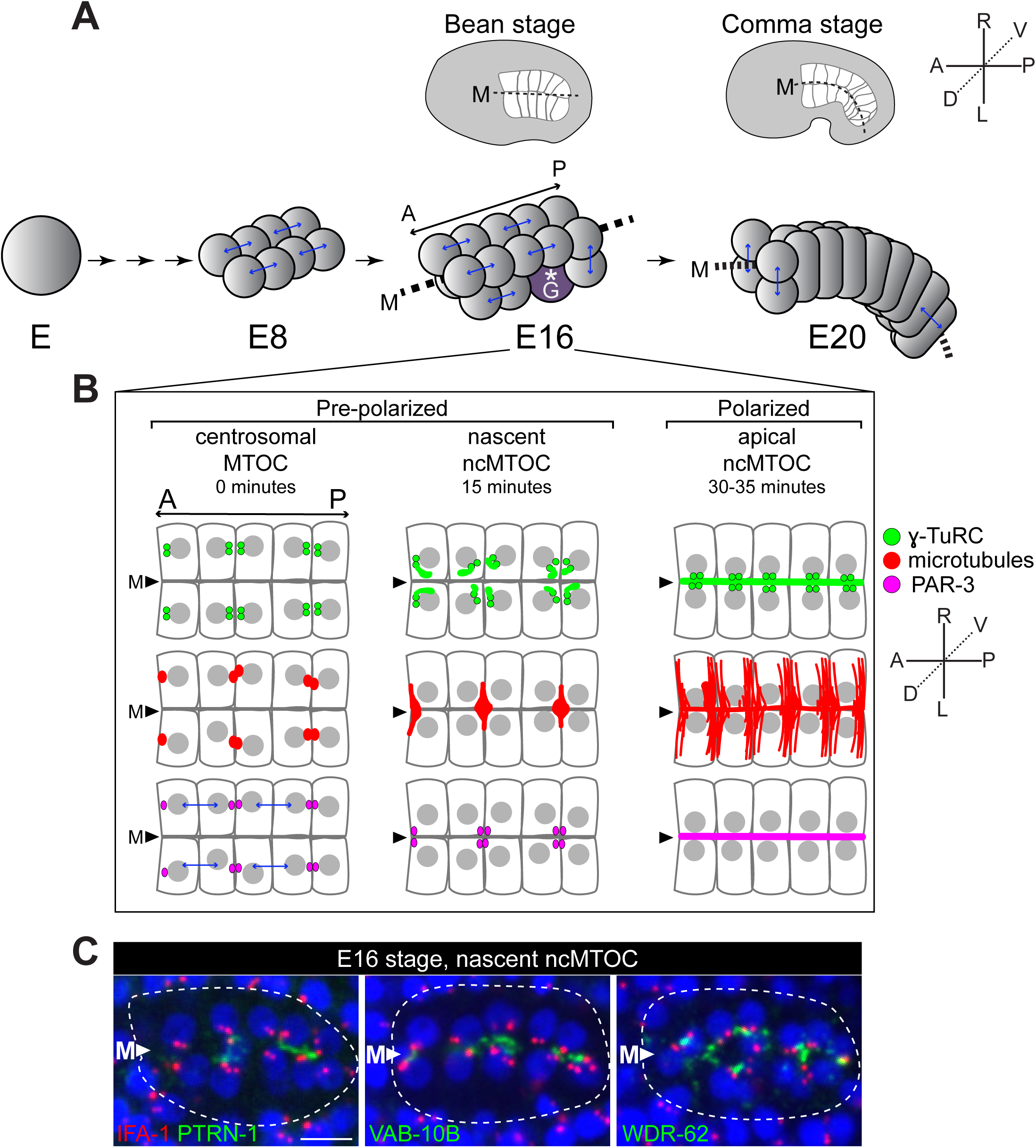
C. elegans intestinal development: A) Cartoon of intestinal development in the *C. elegans* embryo. The intestine is derived from a single “E” blastomere in the embryo. The E cell undergoes four rounds of cell division to give rise to 16 intestinal cells (E16) arranged in two tiers around a central midline (‘M’) at the “bean stage” stage of the embryo. Unless indicated, images throughout this study are shown at E16 from a dorsal view, highlighting 10 of the 16 intestinal cells. Asterisk marks germ cells (‘G’). Approximately 2 hours later (Sulston et al. 1983) as the embryo continues to elongate and the intestine curves, four intestinal cells undergo one more round of division to form the final 20 cells (E20) at “comma stage” of embryogenesis. B) Cartoon showing localization patterns during intestinal polarization. At the exit of the E8-E16 division, the centrosome is the primary MTOC and colocalizes microtubules (red) and γ-TuRC (green). Centrosomes move to the lateral membranes of adjacent non-sister cells where the centrosome is coincident with but does not fully colocalize with PAR-3 (magenta). Centrosomes, apical polarity proteins including PAR-3, microtubules, and γ-TuRC then move toward cell vertices at the future apical surface. During this migration, the nascent ncMTOC is established, thus γ-TuRC exists in two populations: centrosomes (puncta), and a non-centrosomal “plume” that marks the nascent ncMTOC. This reassignment of MTOC function is marked by the inactivation of the centrosome as an MTOC. PAR-3, γ-TuRC, and microtubules then spread across future apical membranes, thereby establishing apical polarity and the apical ncMTOC ∼30 minutes after the E8-E16 division. C) Immunofluorescence imaging of centrosomes (*α*IFA-1, red) relative to PTRN-1::GFP, VAB-10B::ZF::GFP, WDR-62::ZF::GFP (*α*GFP, green) and nuclei (DAPI, blue). Intestine outlined by white dotted line, midline marked with white triangle (‘M’). Note fluorescence intensity is not scaled the same across markers. Scale bar = 5 μm.

**Figure S2.**
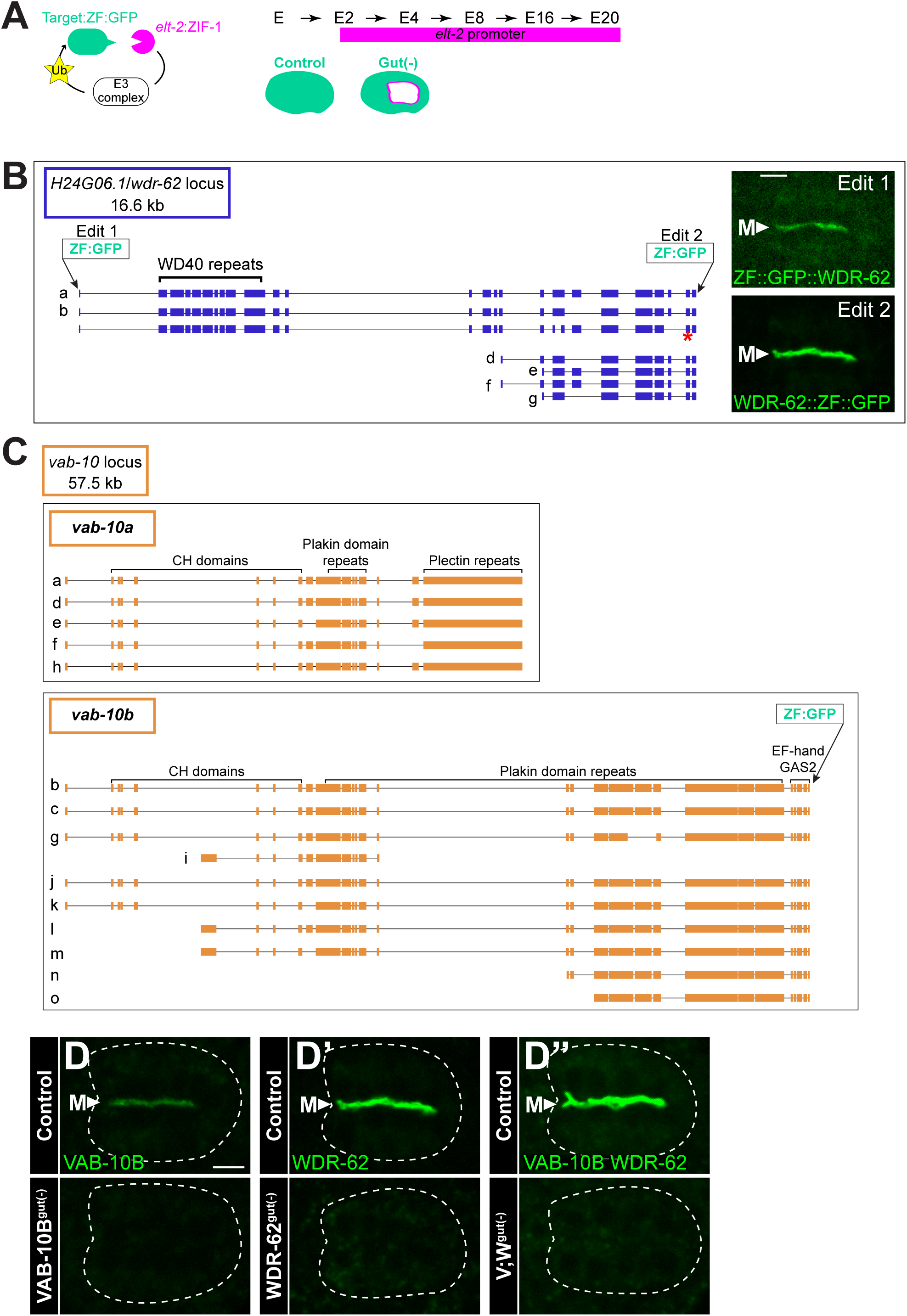
Endogenous tagging and depletion of WDR-62 and VAB-10B. A) Cartoon outlining intestine-specific protein depletion driven by the ZF/ZIF-1 degradation system, a two-component system involving 1) a ‘ZF’ degron tag and 2) the E3 ligase adapter ZIF-1. A protein of interest is endogenously tagged with the ZF tag and GFP using CRISPR/Cas9, and ZIF-1 is over-expressed under an intestine-specific promoter (*elt-2*p). The promoter *elt-2*p is active beginning at the E2-E4 stage of intestinal development and continues driving expression into larval stages. If ZIF-1 is not expressed in the gut, the ZF-tagged protein is not depleted (Control), but intestine-specific ZIF-1 expression drives ZF-tagged protein degradation (‘gut(-)’). B) The *H24G06*.*1/wdr-62* locus was endogenously tagged at indicated positions either at the N-terminus (‘Edit 1’) or C-terminus (‘Edit 2’) with a ZF tag and GFP using CRISPR/Cas9. RNA sequencing data suggests the presence of another isoform containing an early stop (red asterisk). It is unknown whether this isoform is translated, but the C-terminal ZF::GFP insertion we used for our degradation studies would not tag the resulting protein. Live imaging shows GFP localization in polarized intestine from dorsal view for each edit; note fluorescence intensity in images is scaled differently because WDR-62::ZF::GFP is expressed at a higher level than ZF::GFP::WDR-62. C) The *vab-10* locus was endogenously tagged at the indicated locus at the C-terminus with a ZF tag and GFP using CRISPR/Cas9. Domain indications are approximations; for domain analysis see other sources (Bosher et al. 2003; Gally et al. 2016). D-D’) Localization of VAB-10B::ZF::GFP and WDR-62::ZF::GFP in the bean stage polarized intestine without ZIF-1 expression (Control); expression of ZIF-1 in intestinal cells results in tissue-specific degradation of these proteins: VAB-10B^gut(-)^ and WDR-62^gut(-)^, respectively. Images taken at same laser and exposure settings and scaled the same. Dashed white line outlines intestine. (D’’) Expression pattern of VAB-10B::ZF::GFP and WDR-62::ZF::GFP (control) or simultaneous depletion of VAB-10B and WDR-62 (V;W^gut(-)^). Scale bars = 5 μm.

**Figure S3.**
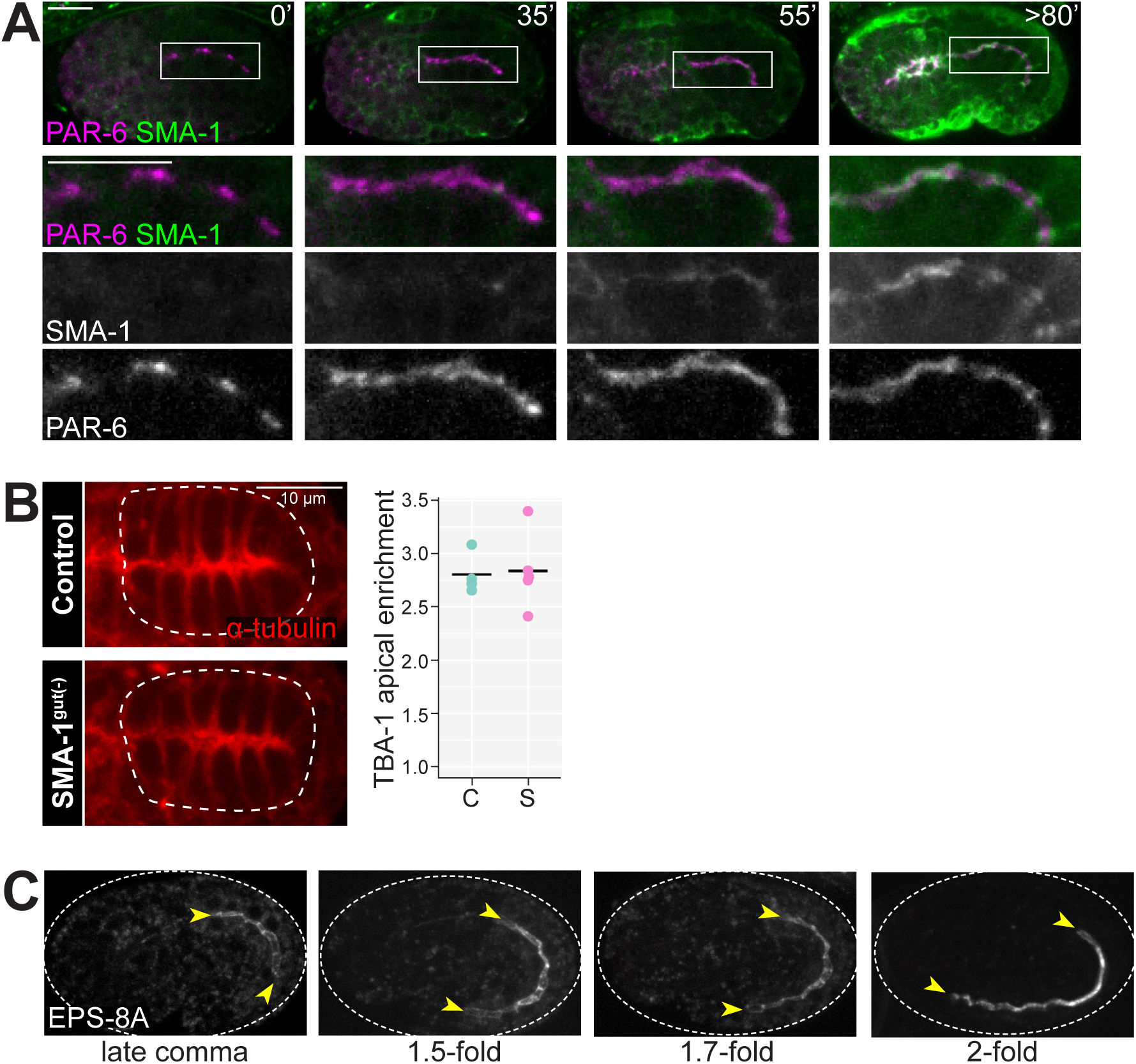
Localization and functional analysis of PTRN-1 proximity interactors: A) Apical polarity marker PAR-6 (magenta) localizes at the apical surfaces of intestinal cells (boxed region, enlargement shown below) before SMA-1 (green). SMA-1 localizes to apical membranes of intestinal cells at late bean/early comma stage of embryonic development. B) Dorsal view of intestine in a live control embryo without protein depletion (‘C’) or with intestine specific depletion of SMA-1 (SMA-1^gut(-)^, ‘S’) expressing mCherry::TBA-1/α-tubulin. Graph on right shows quantification of mCherry::TBA-1 fluorescent signal enrichment at the apical midline for indicated genotypes; not statistically different. C) EPS-8A was first visible in the intestine beginning at the late comma/1.5 fold stage and localized along the apical surface (yellow arrowheads) and increased in expression as embryonic elongation proceeded. Scale bars = 10 μm.

**Figure S4.**
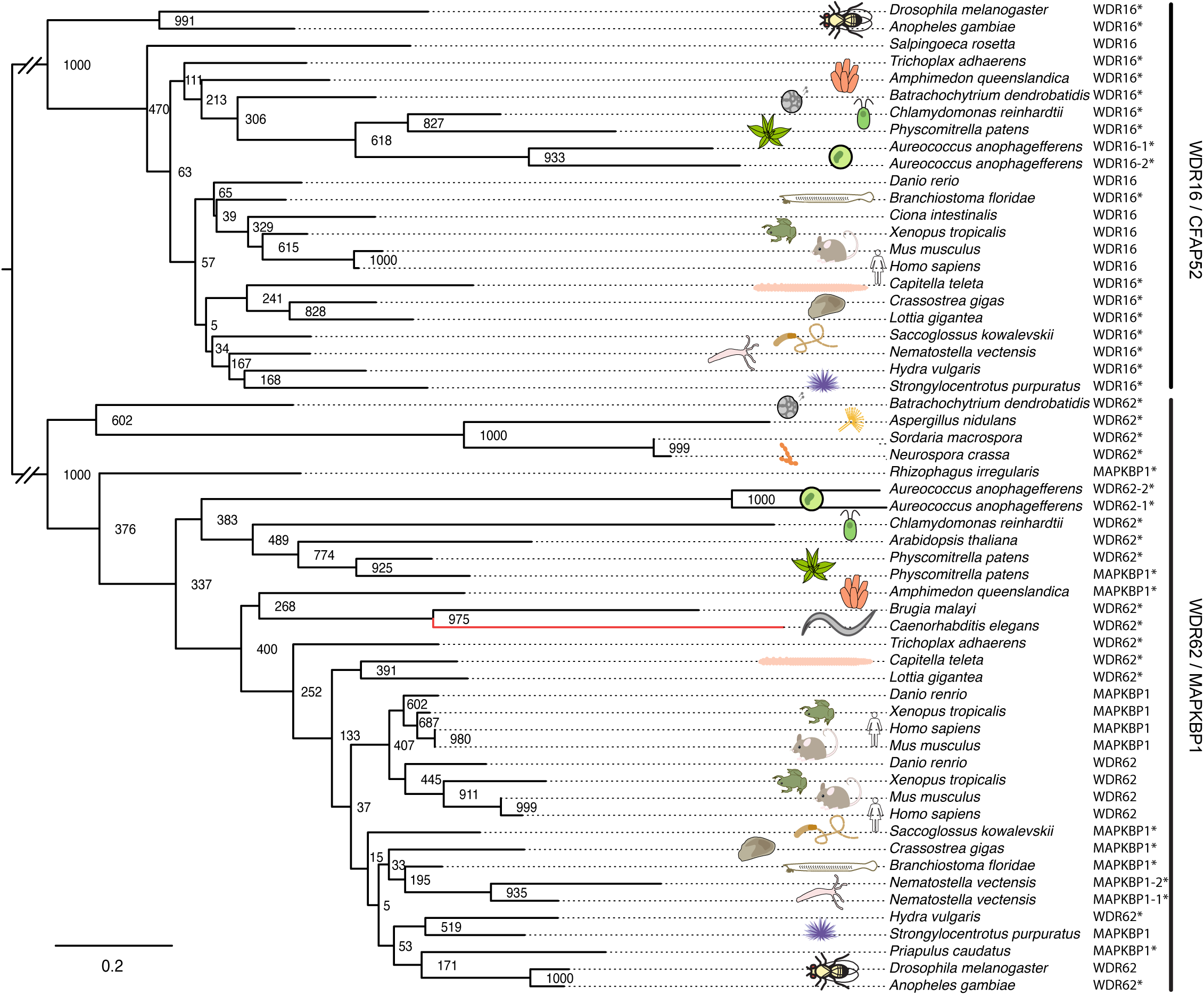
Extended phylogenetic analysis of WDR62 family proteins. *C. elegans* H24G06.1 is a divergent member of the WDR62 protein family of WD40 repeat proteins. Phylogenetic tree of conserved regions within the WD40 repeat region for WDR62 and WDR16 family members (accession numbers available in Table S2) is shown with maximum likelihood support. Double diagonal bars indicate removal of length equal to two scale bars (0.2 substitutions per site).

